# Cas10 residues lining the target RNA binding channel regulate interference by distinguishing cognate target RNA from mismatched targets

**DOI:** 10.1101/2025.04.15.649021

**Authors:** Sarah A. Khweis, Mason Blackburn, Megan O. Pierce, Colby R. Lewis, Jack A. Dunkle

## Abstract

Type III CRISPR systems are defined by the presence of the Cas10 protein and are among the most abundant CRISPR systems in nature. Cas10 forms a complex with crRNA and several Cas proteins that surveils bacterial cells for foreign RNA molecules and when they are detected it activates a cascade of interference activities. The synthesis of the cyclic oligoadenylate signaling molecule by Cas10 is a key aspect of the interference cascade. Despite structures of the Cas10 complex bound to target RNAs, the molecular mechanism by which Cas10 senses the bound state to license interference is lacking. We identified five residues in *S. epidermidis* Cas10, two in the Cas10 Palm2 domain and three in domain 4, that line the target RNA binding channel. We assessed the contribution of these residues to interference in the context of a cognate or mismatched target RNA. We found that the residues regulate whether a mismatched crRNA-target RNA duplex is able to activate interference in vivo. We purified two site-directed mutants of Cas10-Csm and show with in vitro cOA synthesis assays they demonstrate enhanced discrimination of cognate versus mismatched targets.

## Introduction

CRISPR-Cas systems, present in approximately 85% of archaea and 40% of eubacteria, protect these organisms from foreign genetic elements and predation by bacteriophage by detecting the presence of a foreign nucleic acid and mounting an interference response (Barrangou et al. 2007; Makarova et al. 2020). Several steps in this process are common to all CRISPR-Cas systems including crRNA biogenesis, assembly of a ribonucleoprotein complex containing a crRNA bound to a Cas effector protein and surveillance of the cell for nucleic acids that can base-pair to the crRNA which activates interference (Nussenzweig and Marraffini 2020). However, important differences also exist. Some systems sense foreign DNA while others sense RNA and the architecture of the crRNA-bound effector complex varies substantially.

CRISPR-Cas systems have been grouped into two classes and six types (Makarova et al. 2015). Classes 1 and 2 are distinguished by whether the crRNA-bound effector is a multi-protein complex (for example the *S. epidermidis* Cas10-Csm complex consists of five distinct polypeptides) or a single protein (Cas9) (Jinek et al. 2012; Hatoum-Aslan et al. 2013). Within each class, systems are further divided into types based on the presence of a signature effector protein (Cas10 in type III systems), and then into subtypes determined by additional signature proteins (Makarova et al. 2020). Therefore types III-A, III-B and III-D belong to class 1 because they are all multi-protein effector complexes and belong to type III because each possess Cas10. They differ at the subtype level because they possess subtle differences in other members of the protein complex such as diverged Cas11 subunits. In III-A, III-B, and III-D systems, a Cas7 multimer holds the bound crRNA and exposes it to the solvent (Makarova et al. 2017). These complexes recognize foreign RNA that can base-pair with the crRNA. The base-pairing event stimulates synthesis of second messenger molecules, cyclic oligoadenylates (cOA), and these molecules bind to ancillary proteins which contribute to interference via RNase, DNase or other activities (Kazlauskiene et al. 2017; Niewoehner et al. 2017; Athukoralage and White 2021; Rouillon et al. 2022; Steens et al. 2024).

The *S. epidermidis* Cas10-Csm complex (SeCas10-Csm), a type III-A CRISPR system, typifies these features. SeCas10-Csm is an approximately 276-318 kDa complex formed of Cas10, Csm2 (Cas11), Csm3 (Cas7), Csm4 (Cas5) and Csm5 bound to a crRNA of length 37 or 43 nucleotides (Hatoum-Aslan et al. 2013; Paraan et al. 2023). The crRNA interacts extensively with the Csm3, Csm4 and Csm5 proteins. Target RNA, that is RNA with complementarity to the crRNA, can form a duplex with it. The Cas10 and Csm2 proteins extensively interact with the bound target RNA and substantial conformational changes occur in the complex upon target binding (You et al. 2019; Sridhara et al. 2022; Paraan et al. 2023). CA_6_, a cOA made of six adenosine monophosphate subunits, is synthesized by Cas10 and this second messenger binds to Csm6 activating its latent RNase activity (Foster et al. 2018; Nasef et al. 2019). This activity provides potent interference against foreign plasmids and bacteriophage (Hatoum-Aslan et al. 2014; Jiang et al. 2016; Johnson et al. 2024)(Fig 1).

**Figure 1.**
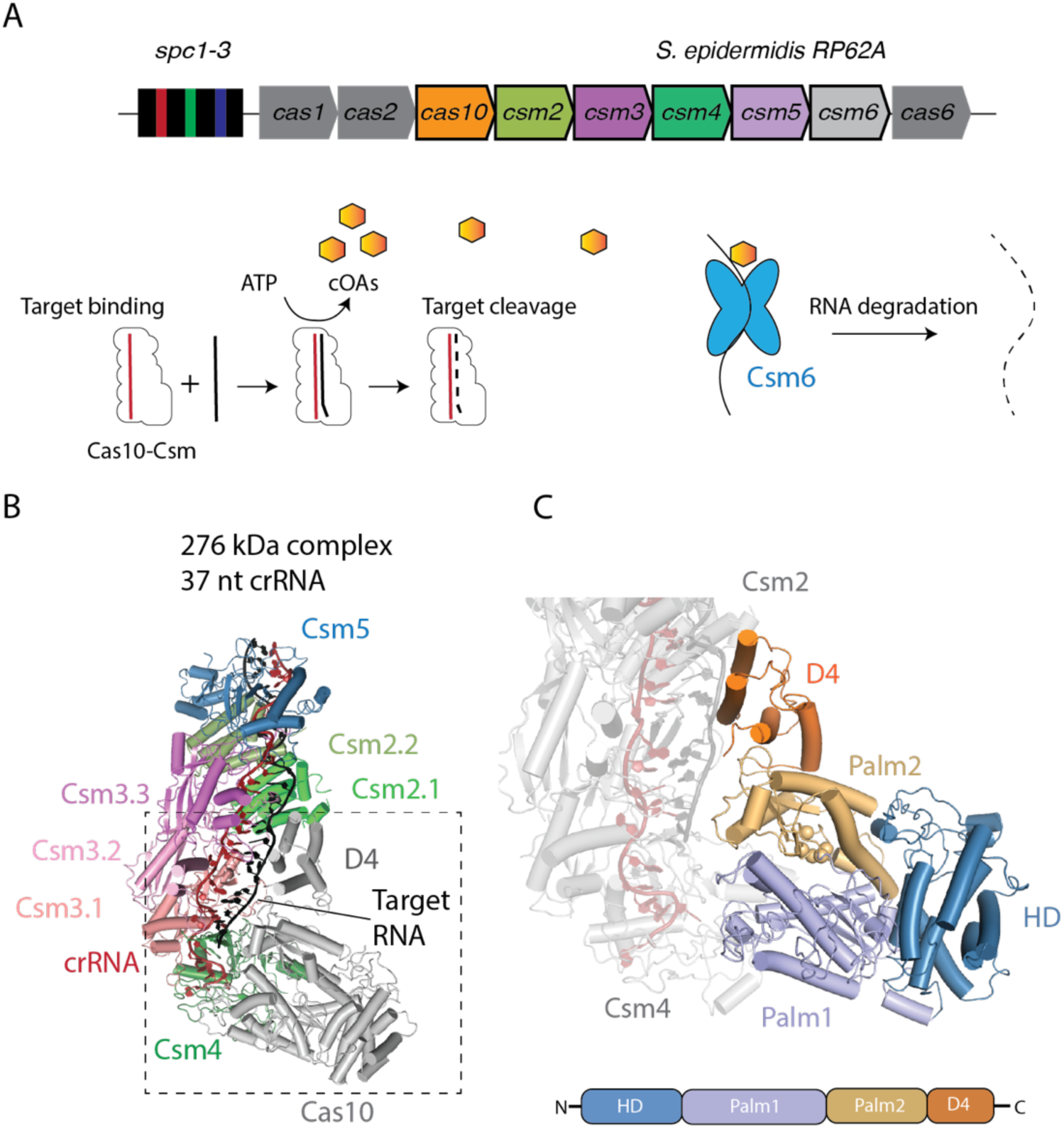
The structure and activity of the Cas10-Csm complex. (A) The CRISPR-Cas10 locus of *Staphylococcus epidermidis* strain RP62A contains three spacers which after transcription and maturation by the Cas6 RNase become crRNAs. Each crRNA associates with Cas10 and Csm2-5 proteins to form the Cas10-Csm ribonucleoprotein complex. This complex can detect foreign RNAs which activates cyclic oligoadenylate (cOA) synthesis. COAs bind to Csm6 and elicit interference by degrading RNA. (B) The Cas10-Csm complex contains a single copy of Cas10, Csm4 and Csm5. In vivo a 276 kDa complex is formed containing three Csm3 proteins, two Csm2 proteins and a 37 nt crRNA. A 318 kDa complex also forms containing a 43 nt crRNA and an extra copy of Csm2 and Csm3. (C) Cas10 is composed of an HD domain which possesses single-stranded DNase activity in many Cas10, a Palm-polymerase like domain—Palm1—fused to a Zn-finger containing region, a Palm2 domain that possesses the active site for cOA synthesis and the C-terminal domain 4 (D4).

The biological role of Cas10-Csm is to detect the presence of foreign RNAs with sensitivity and specificity and couple this detection event to molecular cascades to block replication of foreign nucleic acids. Therefore, molecular diagnostic assays can be built upon the Cas10-Csm complex by simply programming it with a crRNA with complementarity to a pathogen RNA and coupling the molecular cascade to a desirable chemistry for read-out (Gruschow et al. 2021; Santiago-Frangos et al. 2021; Sridhara et al. 2021; Steens et al. 2021). For example, Nemudraia and colleagues showed that the intrinsic signal amplification of Cas10-Csm could be leveraged to detect SARS-CoV-2 RNA at femtomolar sensitivity, equivalent to lateral flow strip diagnostics for SARS-CoV-2, without the use of specialized antibodies or PCR amplification (Nemudraia et al. 2022).

Since both the cOA synthesis activity of Cas10-Csm and its detailed structure were only recently described, a fundamental description of how the complex senses bound target RNA to activate cOA synthesis is lacking (Kazlauskiene et al. 2017; Niewoehner et al. 2017; Jia et al. 2018; You et al. 2019). To address this knowledge gap, we identified two residues in the Palm2 domain of Cas10 and three residues in domain 4 of Cas10 that line the target RNA binding channel and which we hypothesized would influence its interaction with target. We investigated how site-directed mutants of these residues affect interference in cells exposed to cognate and partially mismatched target RNAs. We purified a subset of these site-directed mutants and characterized their target RNA binding and cOA synthesis activity in vitro. We find both in vivo and in vitro that Cas10 residues lining the target RNA binding channel influence the specificity of interference.

## Results

We sought to answer how interference would be affected by the alteration of semi-conserved to conserved Cas10 residues lining the target binding channel which likely engage in non-covalent interactions with target RNA. We believe answering this question will contribute to understanding the mechanistic basis for how cOA synthesis is activated upon target RNA binding to Cas10-Csm. Additionally, we believe this data will be relevant to generating designer Cas10 proteins for diagnostics that are either more tolerant or less tolerant to mismatches in the crRNA-target duplex.

We previously reported a cryo-EM reconstruction of *S. epidermidis* Cas10-Csm bound to an intact target RNA (Paraan et al. 2023). We used this model to identify Cas10 residues which may contact target RNA. The atomic model (PDB code 8do6) contains crRNA, target RNA and proteins Csm2-Csm5. Clear density for Cas10 exists but an atomic model was not deposited due to a lack of clear side chain density. Coordinates for SeCas10-Csm bound to a cleaved target RNA have also been deposited in the PDB, however this structure has undergone substantial conformational changes compared to the structure bound to intact target (Smith et al. 2022). We docked an AlphaFold2 model of Cas10 into the cryo-EM volume we previously reported (EMD-27953) and performed domain-wise rigid body fitting into it (Fig. 2A). We selected Cas10 residues within 6 Å of the target RNA as those with potential non-covalent interactions to the target. The larger than usual distance, 6 Å, was used due to some uncertainty with regard to side chain positions. From the residues identified we noted four basic residues, K524, K628, K691 and R754 which seemed likely to interact with the target (Figs. 2B, 2C). An additional aromatic residue, Y695 from our list of potential interactors was also noted.

**Figure 2.**
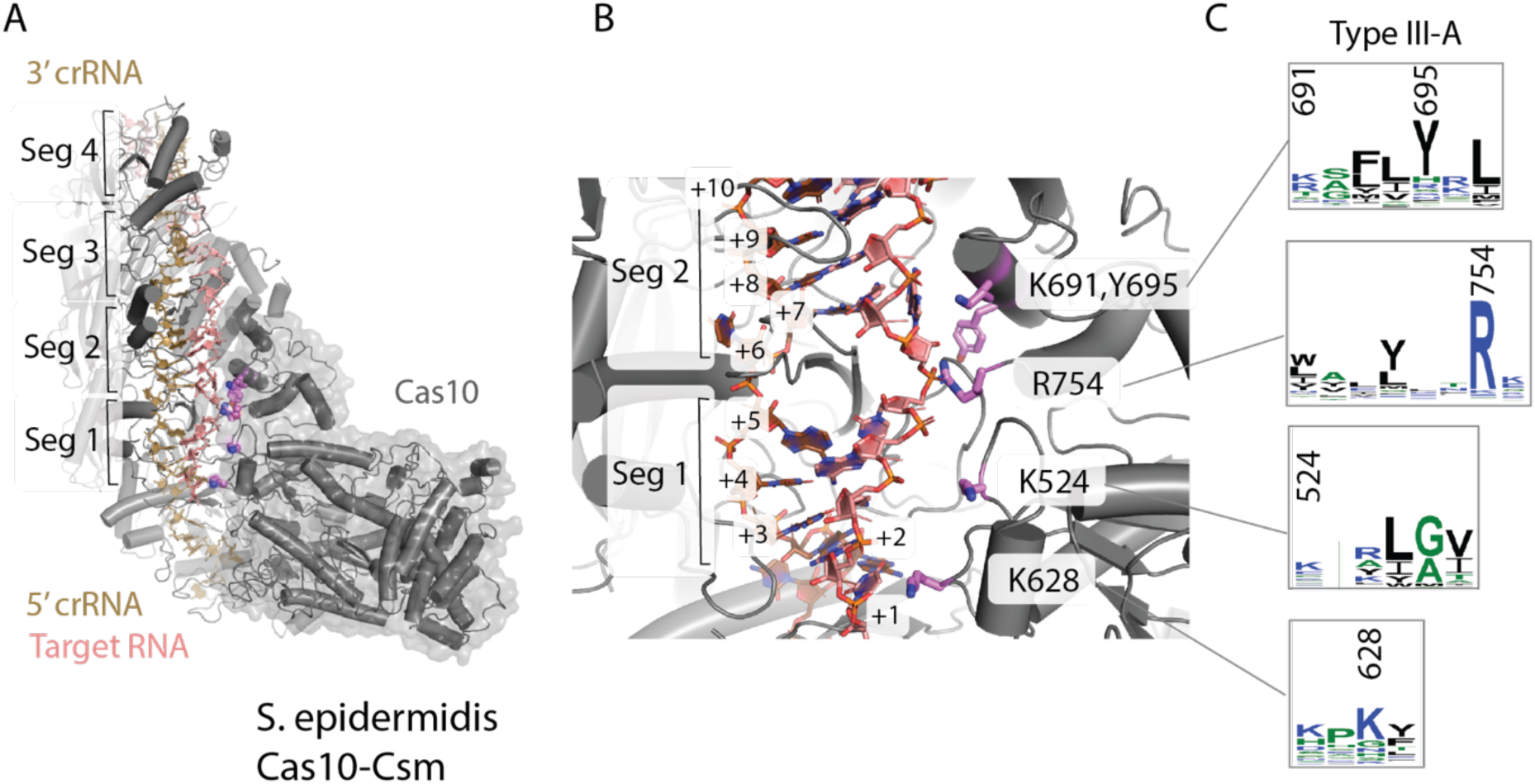
Cas10 in type III-A CRISPR systems contains basic residues and an aromatic residue poised to interact with segments 1 and 2 of the crRNA-target duplex. (A) The structure of *S. epidermidis* Cas10-Csm based on coordinates 8do6 (with an AlphaFold model of Cas10 docked to EM density) is shown. The crRNA-target duplex possesses four segments. Each segment contains five base-pairs and a flipped-out, unstacked nucleotide. (B) Five Cas10 residues poised to contact target RNA are shown as sticks. Cyclic oligoadenylate synthesis by Cas10 is sensitive to mispairing in segments 1 and 2. (C) The Cas10 residues of interest range from moderately to highly conserved in type III-A CRISPR systems.

Several model systems have been reported which assay interference mediated by *S. epidermidis CRISPR-cas10*. This CRISPR-Cas system provides both anti-phage and anti-plasmid immunity. The latter has been assayed in *S. epidermidis* cells utilizing a conjugative plasmid. An extensive set of site-directed mutants and knockouts revealed which *CRISPR-cas10* components were necessary for anti-plasmid immunity specifically revealing that both the Palm2 domain of Cas10, which synthesizes cOA, and the Csm6 nuclease, which is activated by cOA, are required (Hatoum-Aslan et al. 2014). This demonstrates the crucial role of cOA in anti-plasmid immunity. *S. aureus* cells hosting *S. epidermidis crispr-cas10* on a plasmid have also been used to measure anti-plasmid immunity (Pyenson et al. 2017). In these experiments a second plasmid encoding a transcript derived from the *cn20* phage gene, which is complementary to the *spc2* spacer, was used to transform the cells and transformation efficiency was measured. A third model system for anti-plasmid immunity uses *E. coli* hosting a plasmid expressing S*. epidermidis* Cas10-Csm which is then challenged by transformation with a plasmid expressing a *Nes* target (complementary to *spc1*) or a *Cn20* target (complementary to *spc2*) (Fig. 3A). The *E. coli* based system indicates the five proteins that make up the Cas10-Csm complex, a crRNA matured by Cas6 and the Csm6 nuclease are sufficient for anti-plasmid immunity (Foster et al. 2018).

**Figure 3.**
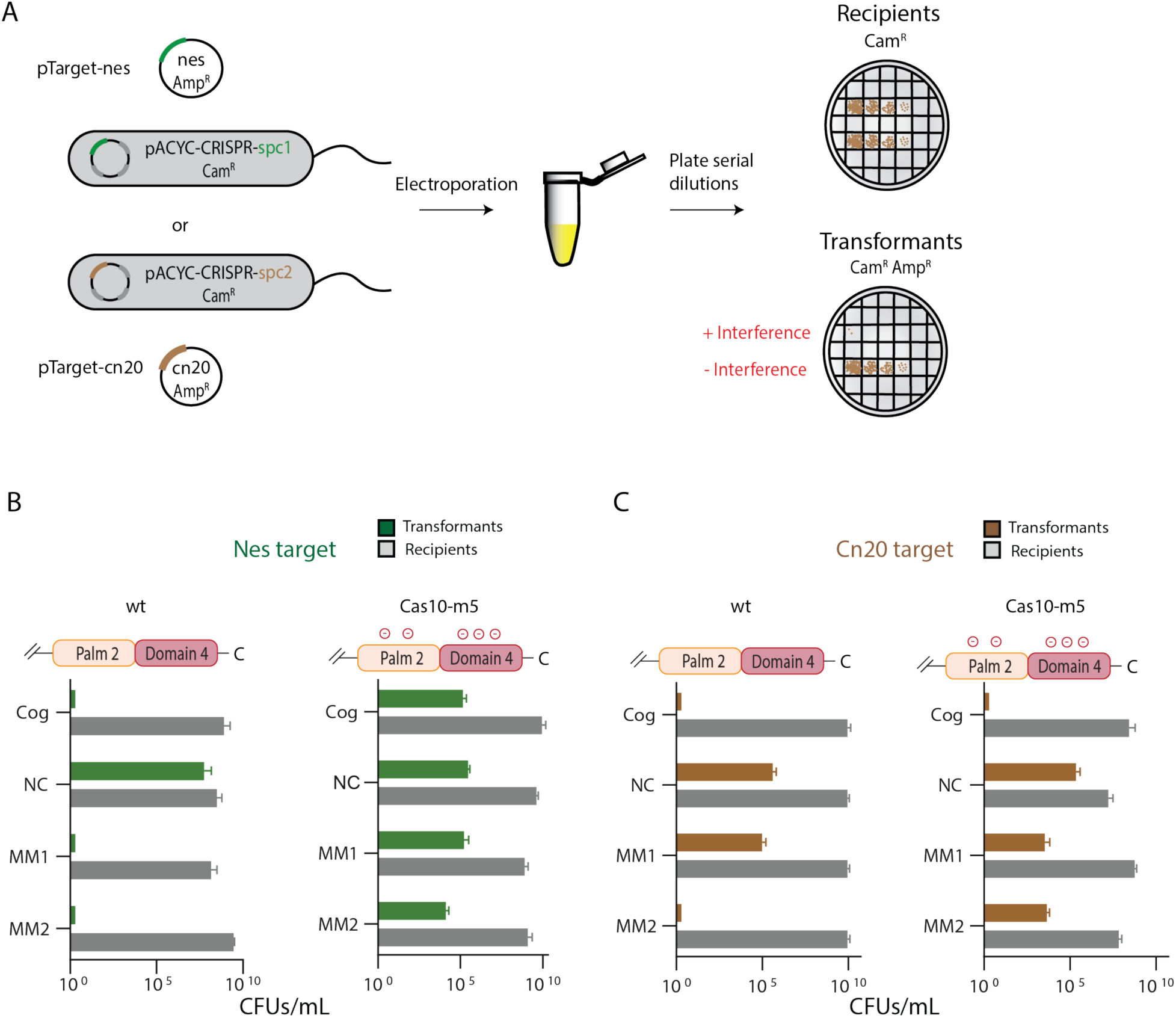
Contacts between Palm2 and domain 4 residues and target RNA contribute to cOA-mediated interference. (A) Diagram of the cOA-mediated interference assay. *E. coli* cells harboring pACYC-CRISPR containing a spacer (*spc1*) that targets the *nes* gene are transformed with pTarget-nes and transformation efficiency reports on whether interference occurred. Alternatively, pACYC-CRISPR containing a spacer (*spc2*) that targets the *cn20* gene is used. (B) Wild type Cas10 interferes with a cognate (Cog) *Nes* target, an *Nes* target in which segment 1 is mismatched (MM1) with crRNA and an *Nes* target in which segment 2 is mismatched (MM2) with crRNA. Cas10 does not interfere with a non-complementary target (NC) which serves as a negative control. A site-directed mutant of Cas10 missing contacts to target RNA in Palm2 and domain 4 (Cas10-m5) does not interfere with the *Nes* targets assayed. (C) Wild type Cas10 interferes with a cognate *Cn20* target and a segment 2 mismatched target but not a segment 1 mismatched target. Cas10-m5 does not interfere with either *Cn20* targets containing a mismatched segment.

We validated the *E. coli* based interference system with extensive controls. In type III CRISPR systems, the crRNA is composed of two sections, the 5’-flank and the body. A target RNA that base-pairs throughout the body section to the crRNA but not with the 5’-flank is termed a cognate target and elicits interference. As a positive control in our assay, *E. coli* expressing wild type Cas10-Csm were transformed with a plasmid encoding a cognate target RNA and as a negative control these cells were transformed with a plasmid encoding a non-complementary target RNA. In each case, the expected results were observed: no transformants were detected in the cognate case and approximately 10^5^-10^7^ colony forming units per mL (CFU/mL) were observed in the non-complementary case (Fig. 3B). In vivo, disruption of Watson-Crick base-pairing at several contiguous positions in the crRNA-target duplex leads to a loss of interference (Maniv et al. 2016; Pyenson et al. 2017; Nasef et al. 2022). We refer to these targets as mismatched. We generated targets that were mismatched at segment 1 (MM1) or segment 2 (MM2) and did this for both the *Nes* transcript and the *Cn20* transcript. When cells expressing wild type Cas10-Csm were challenged with the mismatched targets interference followed a pattern that is consistent with previous data (Nasef et al. 2022). That is, interference was unaffected by the mismatches in the *Nes* sequence context (Fig. 3B). However, in the *Cn20* sequence context MM1 caused a loss of interference but MM2 did not (Fig. 3C). These results confirm previous reports that when the effects of mismatched nucleotides on interference are assayed the identity of the mismatched and the surrounding nucleotides matter (Nasef et al. 2022).

We created a site-directed mutant of Cas10 altered at the five positions described above: K524E, K628E, K691E, Y695E and R754E. We refer to the construct possessing these mutants as Cas10-m5. We utilized the *E. coli* based anti-plasmid immunity system to assay the effect of Cas10-m5 on interference. When presented an *Nes* target, Cas10-m5 is defective in interference for all versions of the target where interference could be expected: cognate, MM1 and MM2 targets (Fig. 3B). When presented a *Cn20* target, Cas10-m5 supports interference on a cognate target but not MM1 or MM2 (Fig. 3C). Therefore, in the *Cn20* context, the mutant residues sensitize Cas10-Csm to mismatches in segment 2 (Fig. 2A). In the *Nes* context, the mutant residues lead to a loss of interference for all versions of this target.

The K524 and K628 residues are located in the Palm2 domain of Cas10 while K691, Y695 and R745 are located in domain 4. To score the relative importance of the two domains in contributing to interference, we next performed interference assays with constructs containing only the two Palm2 mutants (Cas10-mPalm2) or the three domain 4 mutants (Cas10-mDomain4). In the *Nes* context, interference was restored in the presence of cognate target for both Cas10-mPalm2 and Cas10-mDomain4 (Figs. 4A, 4B). Therefore, both Cas10 mutants were more capable of interference than Cas10-m5. In the *Cn20* context, the interference phenotype of Cas10-mPalm2 and Cas10-mDomain4 was similar to Cas10-m5 (Figs. 4C, 4D). Interference occurred on the cognate target but was defective on the mismatched targets. The behavior with the MM1 target was expected. Since wild type is defective in interference with MM1 targets these assays function as a negative control. Notably, transformation efficiency was lower for the MM2 target in these experiments (10^3^ for both Cas10-mPalm2 and Cas10-mDomain4 versus 10^4^ for Cas10-m5) than in the corresponding Cas10-m5 experiment, indicating Cas10-m5 is slightly more defective in interference in this context. In summary, the defects in interference in Cas10-m5 can’t be assigned solely to site-directed mutants of Palm2 or domain 4 but instead both contribute to the property.

**Figure 4.**
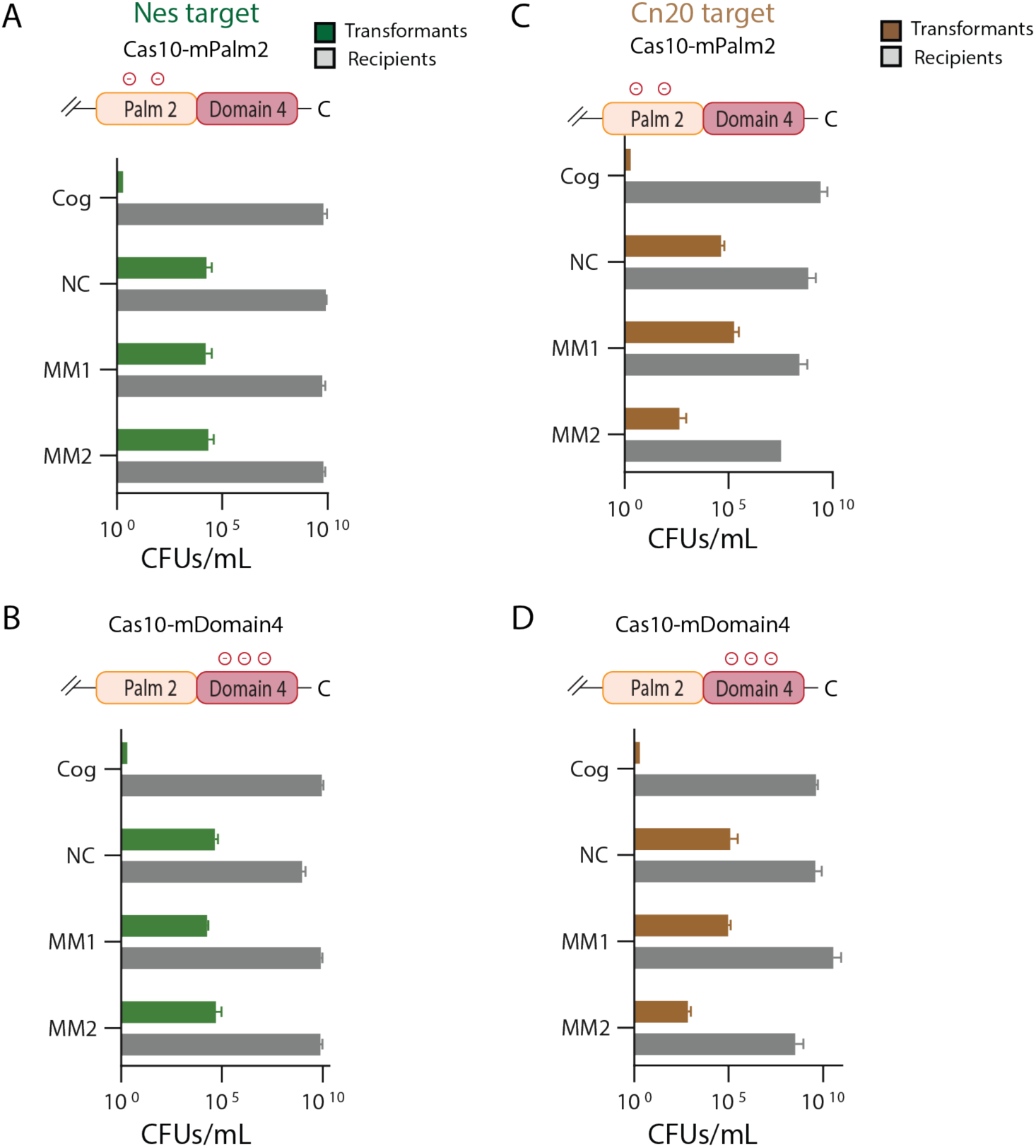
Contacts of either Palm2 to target RNA or domain 4 are sufficient for interference against cognate target RNA but not mismatched target RNA. (A) A site-directed mutant of Cas10 missing contacts to target RNA in Palm2 (Cas10-mPalm2) maintains interference against a cognate (Cog) *Nes* target but loses interference if segment 1 is mismatched to crRNA (MM1) or if segment 2 is mismatched to crRNA (MM2). A negative control, non-complementary target is indicated by NC. (B) A site-directed mutant of Cas10 missing contacts to target RNA in domain 4 (Cas10-mDomain4) was assayed against the panel of targets. (C) and (D) Cas10-mPalm2 and Cas10-mDomain4 were assayed for interference against a panel of targets that contained a *Cn20* target sequence.

We next assayed the contribution to interference of single site-directed mutants of each of the chosen residues. In the *Nes* context, K628E supports interference on cognate, MM1 and MM2 targets behaving identically to wild type Cas10 (Fig. 5A). Interestingly, K524E, K691E, Y695E and R754E support interference with both cognate and MM1 targets but are defective on MM2 targets (Figs. 5A, 5B). These results suggest the four mutant residues each sensitize Cas10-Csm to mismatches in segment 2: while wild type Cas10-Csm possesses a normal interference phenotype on an *Nes* target with MM2, the four single site-directed mutants lose interference. Therefore, the four residues are candidates for a designer Cas10 with enhanced discrimination of mismatches.

**Figure 5.**
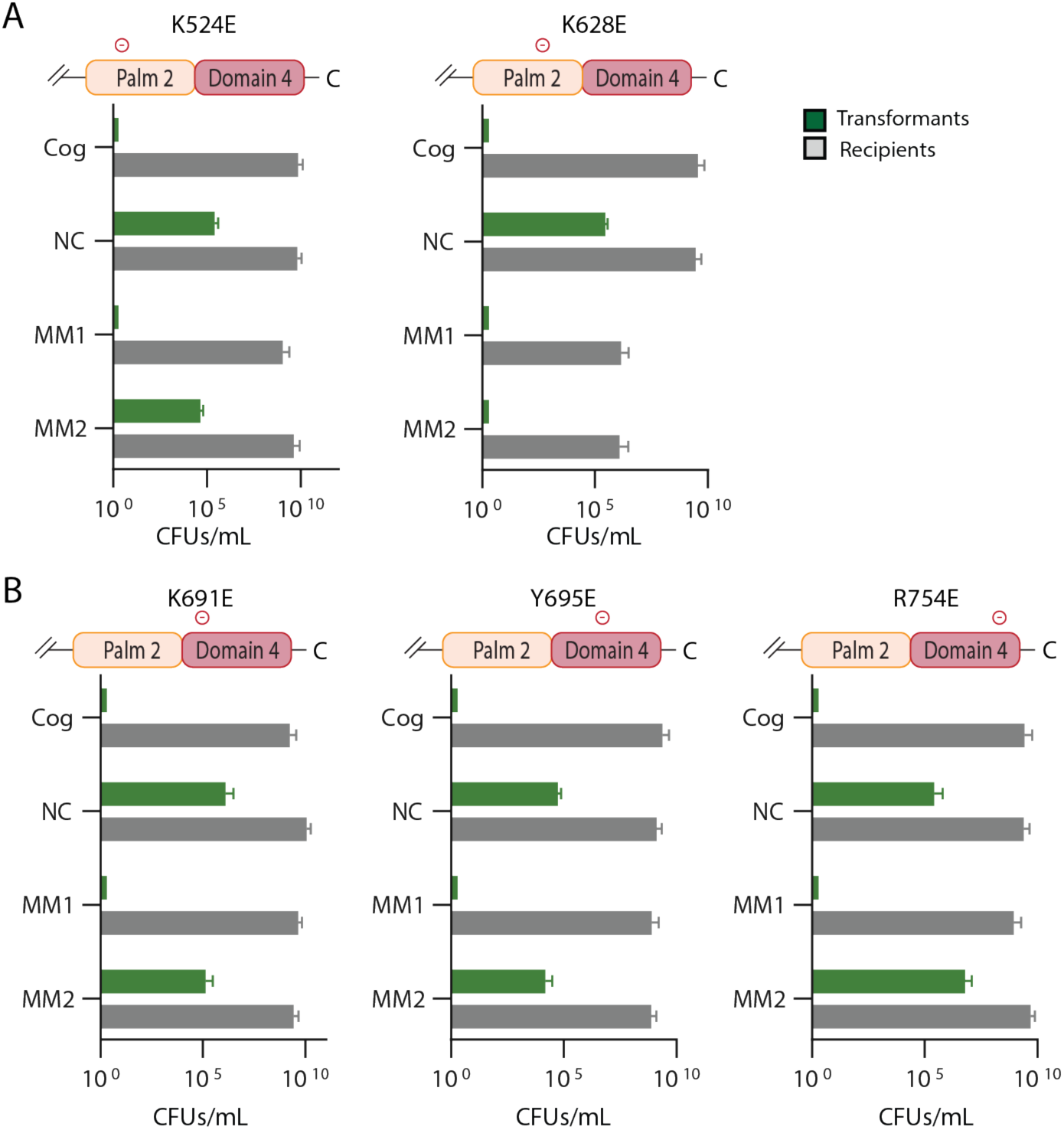
Single site-directed mutant of Palm2 or domain 4 possess interference defects with segment 2 mismatched targets but not with cognate targets. (A) The K524E and K628E site-directed mutants of Palm2 were assayed. (B) The K691E, Y695E and R754E site-directed mutants of domain 4 were assayed. In all cases an *Nes* target was used.

We performed the same assay, measuring interference with single site-directed mutants of Cas10, but now in the *Cn20* sequence context. As noted before, MM1 targets function as a negative control because wild type is deficient in interference with these targets (Fig. 3C). In these experiments, K524E and K691E have interference phenotypes similar to wild type: interference occurs for cognate and MM2 targets. In contrast K628E, Y695E and R754E have a loss of interference for the MM2 target (Figs. 6A, 6B). Considering all the data for single site-directed mutants reveals that K524E, K628E and K691E sensitize Cas10-Csm to mismatches in a sequence context dependent manner while Y695E and R754E sensitize Cas10-Csm to mismatches in both sequence contexts tested (Fig. 7A).

**Figure 6.**
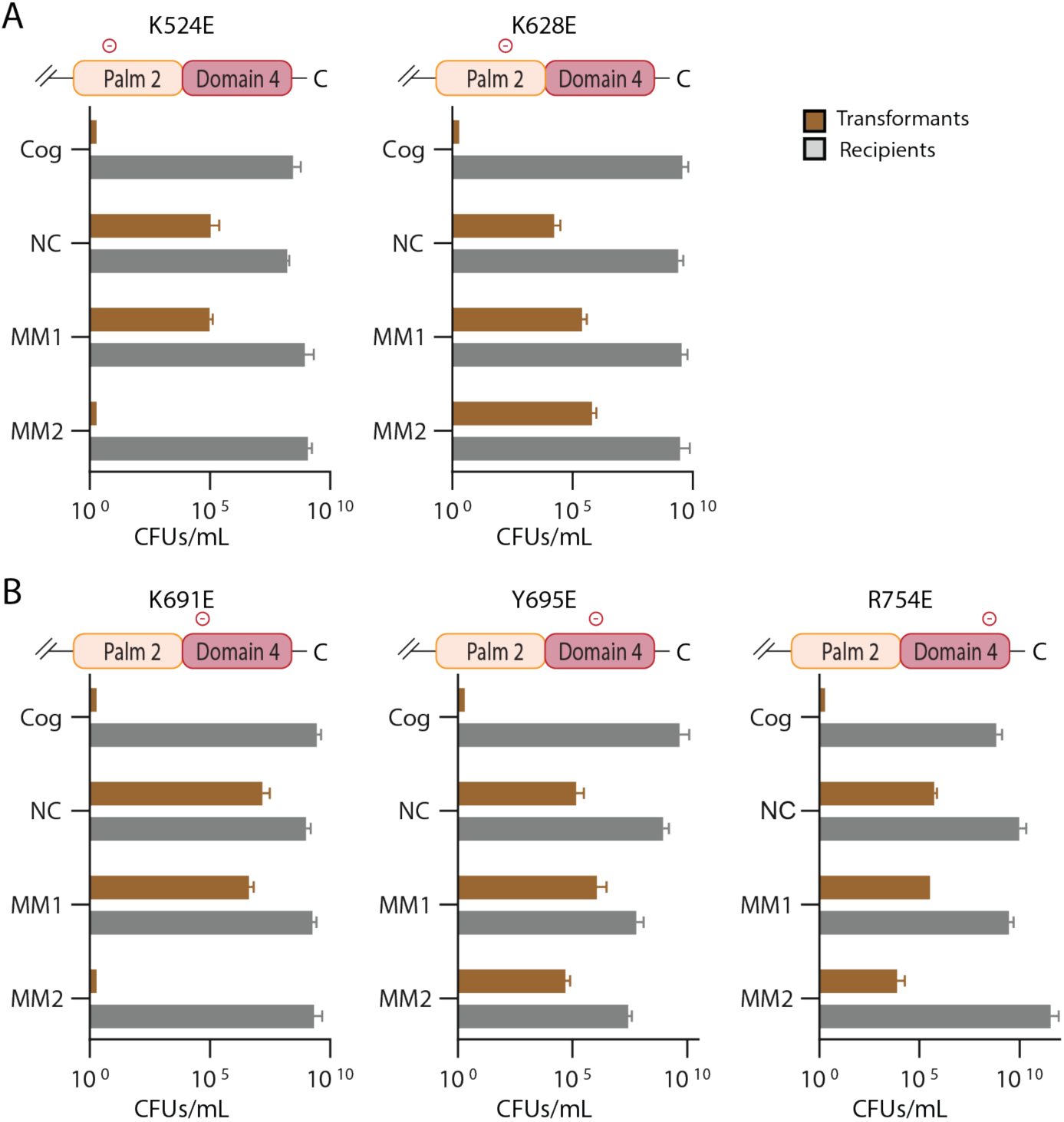
Single site-directed mutants of Palm2 or domain 4 possess interference defects with segment 2 mismatched targets but not with cognate targets. (A) The K524E and K628E site-directed mutants of Palm2 were assayed. (B) The K691E, Y695E and R754E site-directed mutants of domain 4 were assayed. In all cases an *Cn20* target was used.

**Figure 7.**
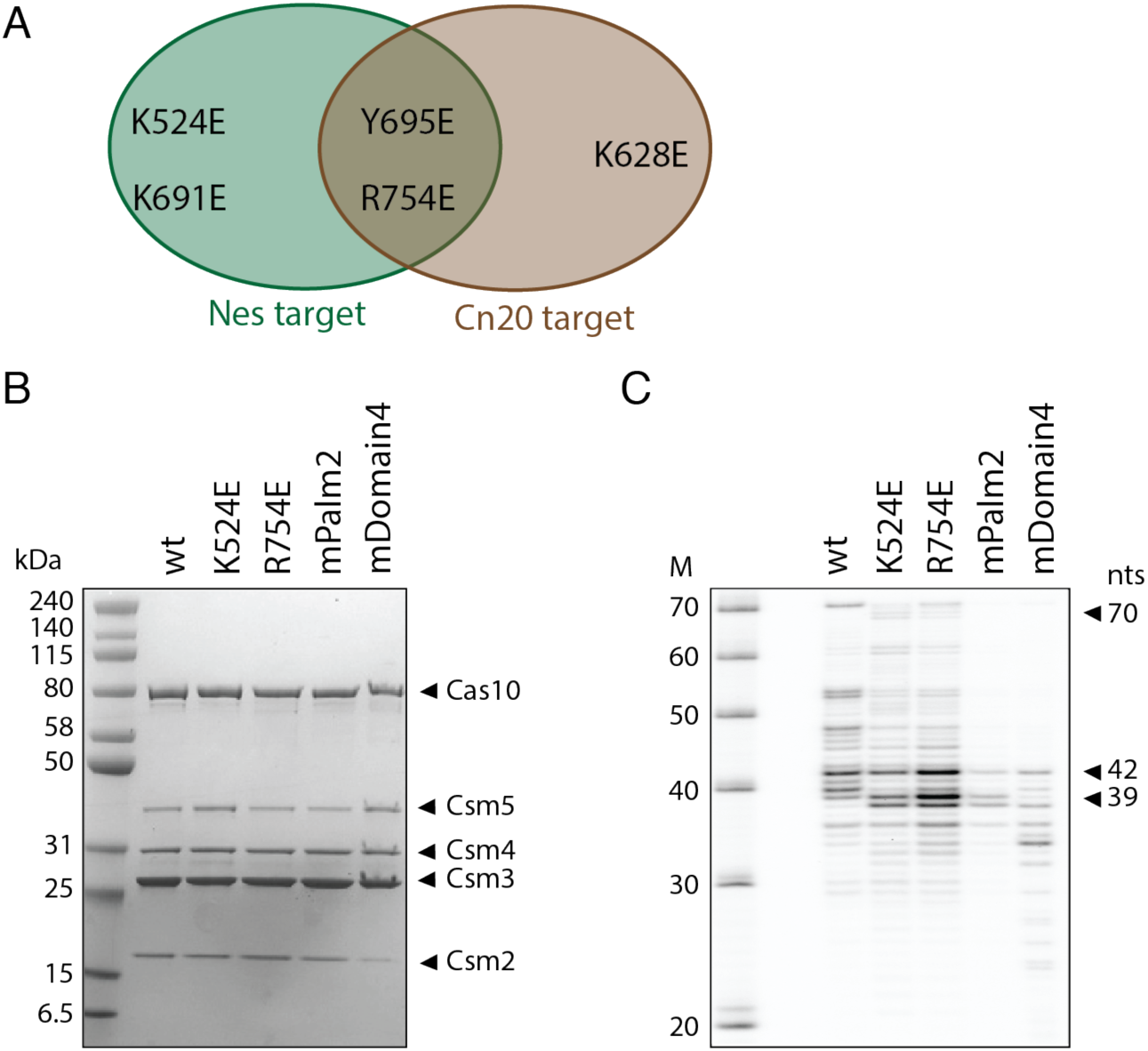
A Venn diagram indicates the scenarios where Cas10 mutants induce sensitivity to mismatches and a PAGE analysis of Cas10-Csm complexes is given. (A) A Venn diagram indicates that Cas10 mutants Y695E and R754E display heightened sensitivity to mismatches in our in vivo interference assay for both *Nes* and *Cn20* targets. (B) SDS-PAGE of site-directed mutants reveals all five Cas10-Csm proteins are present in the purified complexes, however the mDomain4 mutant has diminished Csm2. (C) Urea-PAGE of RNA extracted from site-directed mutants indicates these complexes possess mostly mature crRNA (expected length 37-43 nts) rather than unprocessed, pre-crRNA (length 70 nts). However, the mPalm2 and mDomain4 complexes are depleted in mature crRNA and mDomain4 possesses numerous fragments < 30 nts.

Since Cas10-Csm has utility as a molecular diagnostic in point-of-care settings we next sought to characterize if selected mutants had enhanced discrimination of mismatched targets in vitro. Additionally, in vitro assays allowed us to determine to what degree decreased interference on mismatched targets was due to altered activity versus defects in expression and assembly. We examined how the mutants K524E, R754E, Cas10-mPalm2 and Cas10-mDomain4 affected its behavior in vitro. These four complexes were chosen for two reasons. First, they all support interference in vivo on cognate targets suggesting they may be capable of cOA synthesis in vitro. Second, they are expected to span a range of potential behaviors. For example, K524E sensitizes Cas10-Csm to mismatches in the *Nes* sequence context while R754E sensitizes the complex to mismatches in both sequence contexts tested.

First, we examined whether the four Cas10-Csm mutants could be purified as intact ribonucleoprotein complexes. PAGE analysis indicated all five proteins that comprise the complex are present, although Csm2 appears to be present at sub-stoichiometric levels in mDomain4 (Fig. 7B). We also performed a PAGE analysis of the crRNA content of the mutants. Wild type complex has multiple bands in the range of 29-70 nt with 40 nt and 42 nt being most prominent (Fig. 7C). Notably, there is a greater distribution of crRNA sizes than has been observed when Cas10-Csm is purified from native *S. epidermidis* cells indicating maturation of crRNA during recombinant expression in *E. coli* may proceed via alternate pathways (Walker et al. 2017). However, the 40 nt and 42 nt bands are similar to the most abundant bands, 37 nt and 43 nt, observed when SeCas10-Csm is purified from *S. epidermidis* (Walker et al. 2017). The four mutant Cas10-Csm complexes also possess bands around 40 nt, however the distribution of sizes and crRNA amount differ (Fig. 7C). Analysis of mPalm2 and mDomain4 crRNA suggest they are approximately 42% and 43% respectively as abundant as wild type (Fig. S3). We concluded at this point that the single Cas10 mutants efficiently mature into active Cas10-Csm complexes while the multi-mutants mature with moderate efficiency. We focused the remainder of our in vitro analysis on the single mutants.

Next, we sought to determine whether the K524E and R754E site-directed mutants had normal or altered target RNA binding and importantly how affinity was affected by segment mismatches, MM1 and MM2. To quantitate the effect of the mutants on target RNA binding we performed an affinity binding assay with a fluorescently labeled cognate target RNA and labeled mismatch target RNAs (Fig. 8A). Experiments were performed in the absence of [Mg^2+^] to prohibit target RNA cleavage by the Csm3 component of the complex. Wild type binding to a cognate target RNA served as a positive control and binding to a fluorescently labeled non-complementary RNA (scrambled sequence) served as a negative control. The fraction active for both K524E and R754E was estimated to be approximately equal to wild type. A comparison of the K_d_ values for wild type binding to cognate target compared to K524E and R754E shows that the mutants bind 1.8 and 2.8 fold weaker respectively (Fig. 8B). However, our main interest was in whether the mutants could still bind mismatched targets with high affinity. Compared to cognate targets, K524E and R754E K_d_ values were lower for the MM1 target indicating tighter binding, and were only mildly higher for the MM2 target, 2.7 and 1.6 fold respectively. In sum these site-directed mutants don’t dramatically alter target RNA binding affinity.

**Figure 8.**
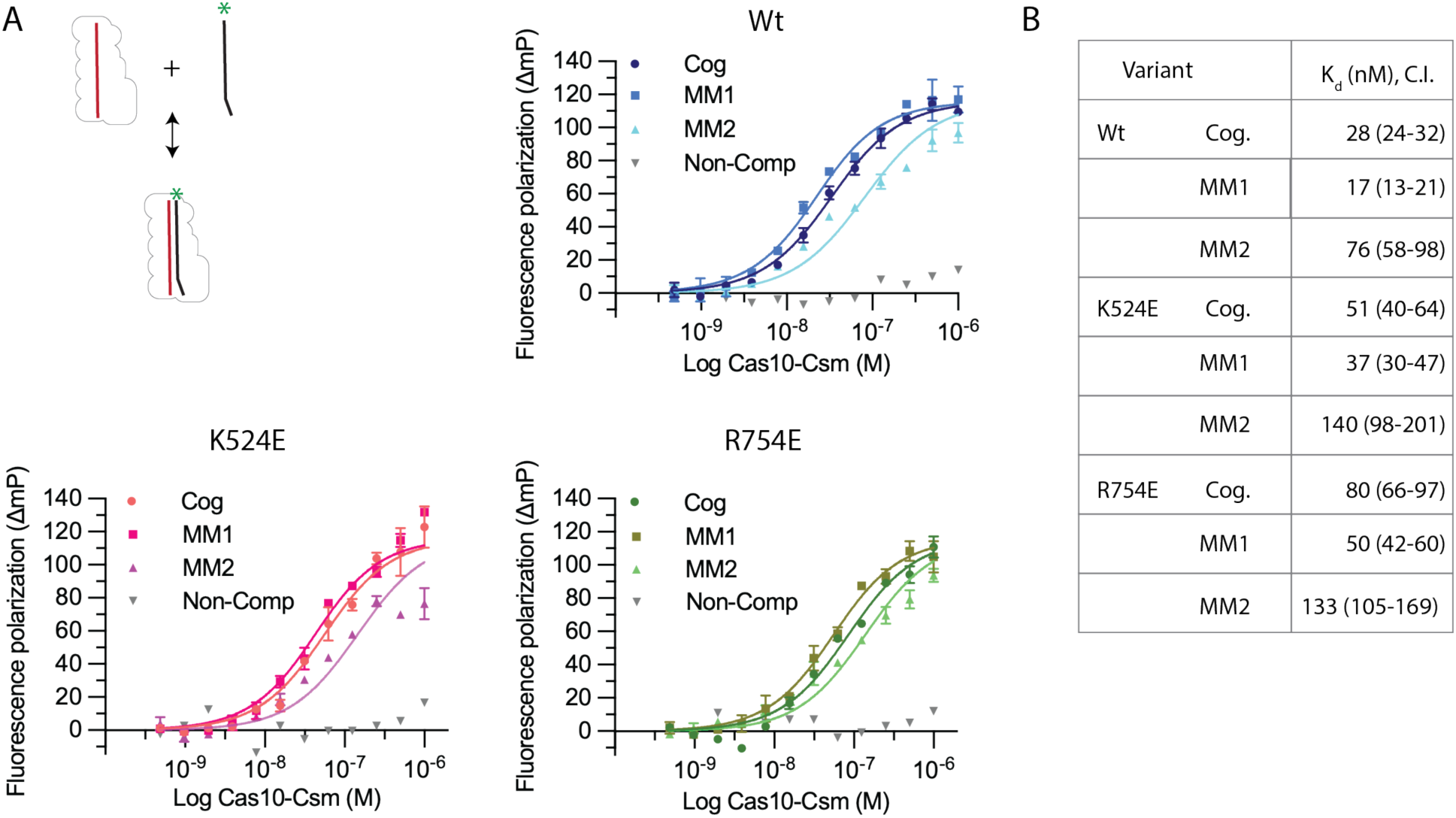
The affinity of target RNA binding to Cas10-Csm variants K524E and R754E. (A) Fluorescently labeled target RNAs were incubated with wild type Cas10-Csm and variants. Binding affinity was measured by fluorescence polarization. Target RNA sequences used are cognate (Cog), a target with segment 1 mismatched (MM1), a target with segment 2 mismatched (MM2) and a non-complementary, negative control target (Non-comp). Two replicate titrations were performed for each target RNA (except the negative control) and best-fit curves were determined by non-linear regression. Mean and standard error are plotted for each point in the titration. (B) Dissociation constant (Kd) values for Cas10-Csm binding to target RNAs. Parentheses indicate 95% confidence interval.

Having confirmed K524E and R754E were competent in target RNA binding we sought to determine how cOA synthesis may be altered. We hypothesized that cOA synthesis by Cas10 site-directed mutants may be more sensitive to mismatches in the crRNA-target duplex than wild type Cas10-Csm. To test this, we in vitro transcribed a panel of *Nes* target RNAs containing either cognate sequence or a single mismatch in positions +1 to +11 of the duplex, which comprise segments 1 and 2. This is similar to boundaries of the crRNA-target duplex described as the Cas10 activating region (Steens et al. 2021). We performed cOA synthesis reactions in vitro with purified Cas10-Csm, a member of the target RNA panel and α-^32^P-ATP. The radioactive cOA products were separated by TLC and quantitated (Figs. 9A, 9B). It has previously been shown that when wild type SeCas10-Csm is presented an *Nes* target with mismatches in either position +2, +5, +7, +8 or +11 a substantial reduction of cOA products occurs (Nasef et al. 2019). We observed that for the K524E and R754E mutants this phenomenon is enhanced. To visualize this, we generated a volcano plot to visualize the statistical significance derived from three replicate reactions plotted against the fold change (Fig. 9C). Notably, fold change is a comparison of the cOA synthesis reaction using a mismatched target to the reaction for the same Cas10 variant but in the presence of a cognate target. Dashed lines highlight the reactions with p < 0.05 that also have pronounced defects in cOA synthesis. Since we have plotted fold change versus cognate target, the reactions left of the vertical dashed line demonstrate that for specific mismatches the K524E and R754E mutants possess enhanced discrimination against mismatched targets. That is, K524E and R754E possess 42% and 49% respectively of the activity on a cognate target exhibited by wild type but have an outsize defect in cOA synthesis on several mismatches in the Cas10 activating region. The positions of greatest interest are labeled in Figure 9C.

**Figure 9.**
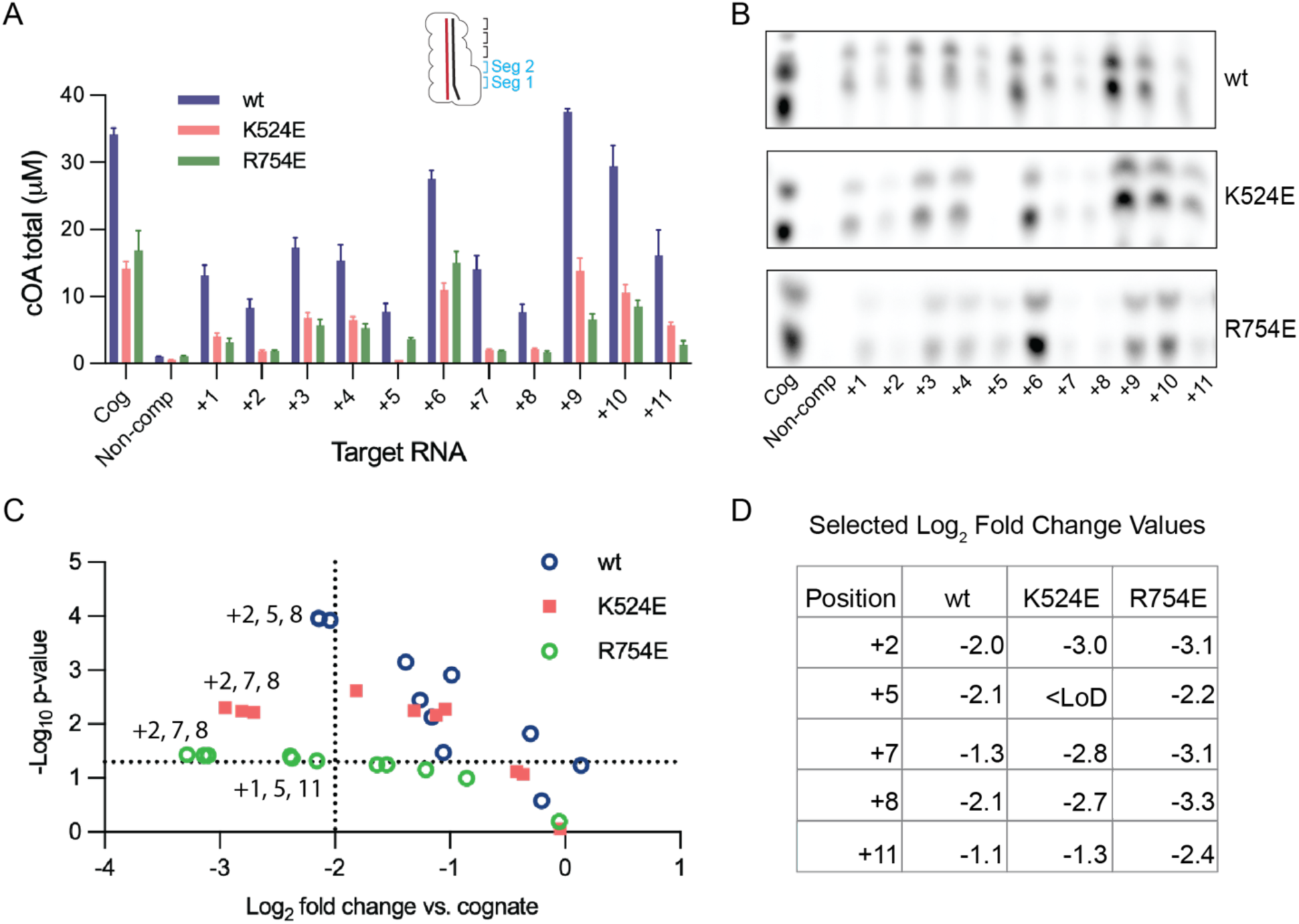
Cas10 mutants enhance sensitivity to single mismatches during in vitro cOA synthesis. (A) Target RNA based on the *Nes* sequence with single mismatches at positions +1 through +11 was used to stimulate cOA synthesis by Cas10-Csm. The amount of cOA produced from three replicate reactions was quantified. The mean and standard error of the mean are plotted. COA amounts for non-complementary target RNA and the K524E variant stimulated by the +5 target RNA were below the limit of detection and are plotted at that limit, y = 1.0 μM. (B) Images of representative TLC plates used for cOA quantitation. (C) A volcano plot of the data in panel A is shown. The amount of cOA synthesized by each Cas10 variant when bound to a single mismatch target RNA is compared to the cOA amount when a cognate target is used. The K524E +5 data point is not plotted because it is below our limit of detection (LoD). (D) A table of selected log_2_ fold change values is given.

## Discussion

Proper licensing of interference in CRISPR-Cas systems is important for bacterial fitness. Overly strict base-pairing requirements between crRNA and targets could lead to the failure of a bacterium to activate interference against a virus that accrued subtle sequence variations since spacer acquisition. Conversely, loose base-pairing requirements could lead to interference being activated by native bacterial genes. For type III-A CRISPR systems the mechanisms of licensing have not been fully described. We performed in vivo and in vitro experiments to determine whether residues lining the target RNA binding channel contribute to Cas10-Csm’s ability to distinguish cognate from mismatched targets and found that they do. Both in vivo and in vitro experiments support this conclusion. When assayed in vivo, the site-directed mutants K524E, K628E and K691E had sequence context specific effects on target discrimination. By contrast, Y695E and R754E enhanced target discrimination in vivo for both sequences, *Nes* and *Cn20*, we tested. In vitro experiments further supported our finding: K524E and particularly R754E displayed enhanced discrimination for some single mismatches between crRNA and target RNA. The in vitro result has bearing on the design and utility of nucleic acid diagnostics based on the Cas10 CRISPR system.

Comparison among our in vivo and in vitro data reveals broad agreement, however, certain specifics require explanation. The mPalm2 and mDomain4 multi-mutants display robust interference against a cognate target in vivo but in vitro mPalm2 cOA synthesis is 20% of cognate and mDomain4 synthesizes so little cOA that it is below our limit of detection (Fig. S5). There are several possible explanations for this. First, we see evidence of reduced stability of purified mPalm2 due to its lower amount of bound crRNA and for mDomain4 its lower level of bound crRNA and its reduced amount of bound Csm2. Another relevant factor is that the level of cOA required for interference in vivo may be very low and in vivo phenomenon such as product inhibition are not in play: in cells cOA are sequestered by binding to CARF domain containing proteins after synthesis while in vitro they are not sequestered and can potentially participate in product inhibition of Cas10. We also note that segment mismatches between crRNA and target were used in our in vivo assay and that multiple mismatches are required in these assays to observe an interference phenotype. However, in our in vitro cOA synthesis assays, single mismatches between the crRNA and target are observed to be capable of dramatically reducing cOA production. These phenomena have been observed by us and others (Maniv et al. 2016; Pyenson et al. 2017; Rouillon et al. 2018; Nasef et al. 2019; Gruschow et al. 2021; Steens et al. 2021; Nasef et al. 2022; Karneyeva et al. 2024). It has been shown that the complementarity between the crRNA and target required for interference in vivo is affected by target abundance (Karneyeva et al. 2024). Therefore, the discrepancy between the complementarity needed for in vivo interference versus in vitro cOA synthesis may be the result of the fact that our interference assay uses a high copy number plasmid driven by a T7 promoter resulting in high expression of the target.

Due to their utility in gene editing, engineering CRISPR-Cas systems for enhanced specificity has been an area of major interest. A comparison of our work with experiments to engineer high fidelity variants of Cas9 points to future tactics that could be used. Early experiments showed that pruning *S. pyogenes* Cas9 contacts to either the target DNA strand or the non-target DNA strand reduced cleavage of off-target sequenced an approach similar to what we have taken with Cas10 (Kleinstiver et al. 2016; Slaymaker et al. 2016). Later single molecule and time resolved cryo-EM experiments of SpCas9 bound to cognate and mismatched targets revealed the importance of multiple checkpoints and concerted conformational changes that proceed DNA cleavage (Sternberg et al. 2015; Bravo et al. 2022). The HNH domain of SpCas9 is dynamic until the complex binds a cognate DNA which promotes formation of the active conformation of HNH which in turn allosterically activates the RuvC nuclease domain for concerted strand cleavage (Sternberg et al. 2015). Two loops, L1 and L2, that link the HNH domain and the RuvC domain interact with the minor groove of the guide RNA-target DNA heteroduplex and are critical for activating SpCas9 along with of kinking of the heteroduplex (Bravo et al. 2022). Recognition of the role of these events in specificity and structures of SpCas9 bound to mismatched targets has led to new approaches for engineering SpCas9 specificity (Bravo et al. 2022; Pacesa et al. 2022). It has been demonstrated that Cas10-Csm undergoes significant conformational changes in Csm2 and Cas10 upon target RNA binding (You et al. 2019; Sridhara et al. 2022; Paraan et al. 2023). A clearer understanding of how these conformational changes specifically promote Cas10 activation may open new avenues for engineering Cas10.

The realization that CRISPR-Cas systems are well-suited to serve as molecular diagnostics was realized early in the history of CRISPR biotechnology (East-Seletsky et al. 2016; Gootenberg et al. 2017). CRISPR-Cas systems have been harnessed to detect RNA viruses, DNA viruses and human single nucleotide polymorphisms (Gootenberg et al. 2017; Chen et al. 2018; Gootenberg et al. 2018; Myhrvold et al. 2018). Each CRISPR-Cas system has intrinsic strengths and limitations as a diagnostic tool. For example, as an RNA targeting system, Cas10 is well-suited for detection of RNA viruses and has been deployed to sensitively detect SARS-CoV-2 RNA (Gruschow et al. 2021; Santiago-Frangos et al. 2021; Sridhara et al. 2021; Steens et al. 2021). A strength for Cas10-Csm is its ability for signal amplification: each target RNA detection event initiates a multiple turnover reaction producing cOA which then activate a CARF domain containing enzyme, such as Csm6, which also performs a multi-turnover reaction such as collateral RNA cleavage. A limitation of Cas10-Csm is its loose specificity. Our data show rational mutagenesis can improve the specificity of Cas10-Csm. An outstanding question is whether directed evolution of Cas10 could also enhance its specificity.

## Materials and methods

### Docking of Cas10 into cryo-EM density

The AlphaFold2 model of *S. epidermidis* Cas10 given by AF-Q5HK89-F1-v3 was docked into the density given by EMD-27593 using real space refinement implemented in Phenix (Liebschner et al. 2019). Each domain of Cas10 was fit as an independent rigid body. The PDB coordinates 8do6 give the atomic model of the Csm2-5 proteins, crRNA and target RNA for this map.

### Site-directed mutagenesis

The pACYC vector encoding *S. epidermidis* CRISPR-Cas10 was a gift from Dr. Michael Terns of the University of Georgia (Ichikawa et al. 2017). The CRISPR-Cas10 locus encodes for a single spacer gene, *spc1*, which targets the *Nes* transcript, denoted pACYC-CRISPR-*spc1*. In a previous study, we developed a vector that replaced the *spc1* gene with *spc2*, resulting in pACYC-CRISPR-spc2 (Nasef et al. 2022). Both pACYC plasmids possess a chloramphenicol resistance marker. Maps of the two plasmids are given in Supplementary Data (Fig. S1) and plasmid sequences and all oligo and gene fragment sequences are also given (Table S1). For the site-directed mutants of pACYC-CRISPR-spc1, Q5 site-directed mutagenesis was used to introduce single, double, and triple mutations (New England Biolabs). For the quintuple mutant, a GeneArt string (ThermoFisher Scientific) was ordered containing the 5 mutations along Cas10. Inverse PCR was used to linearize pACYC-CRISPR and Gibson Assembly was used to ligate the gene string into the vector. To create a pACYC-CRISPR-spc2 plasmid with Cas10 mutants, we began with the pACYC-CRISPR-spc1 plasmids carrying the *cas10* mutants. We utilized NcoI and PspXI restriction sites to remove *Nes* targeting *spc1* and DNA ligase to insert a GeneArt string (ThermoFisher Scientific) encoding *spc2* that targets *Cn20*. All plasmids were verified by whole-plasmid sequencing.

A pTRC vector containing an ampicillin marker and a *nes* target gene (pTarget-nes) was also provide by Dr. Michael Terns. We reported the insertion of a *cn20* target gene, into this plasmid to create pTarget-cn20 in a previous study (Nasef et al. 2022). In our current study, we constructed two new target plasmids: each has mismatches from +7 to +12 along the crRNA-target duplex and is denoted as MM2. Gibson assembly was used to introduce MM2 into pTarget-nes with a gene strand (Eurofins Genomics). For experiments targeting the *cn20* gene, Q5 PCR was used to introduce MM2. All constructs were verified by whole plasmid sequencing. The target plasmids denoted MM1 carrying mismatches in the crRNA-target duplex in positions +1 to +6 were reported previously (Nasef et al. 2022).

### Interference Assays

*E. coli* BL21(DE3) cells containing pACYC-CRISPR wild type plasmids were made electrocompetent then transformed by electroporation with 100 ng of the corresponding pTarget plasmid. After transformation, the cells were resuspended with SOC medium and incubated at 37°C with shaking for 1 hour. Ten-fold serial dilutions were then prepared in LB medium, and the resulting suspensions were plated onto LB-agar plates containing either chloramphenicol (17 µg/mL) or a combination of chloramphenicol (17 µg/mL) and ampicillin (50 µg/mL). After overnight incubation at 37°C, the plates underwent quantification of colony-forming units per mL (CFU/mL) for three independent transformations of each target plasmid.

### Expression and purification of Cas10-Csm variants

One Shot BL21-AI (ThermoFisher Scientific) strains harboring pACYC-CRISPR were grown in LB medium at 18°C with shaking until an OD_600_ of 0.8 was reached. The cultures were induced with 0.2% w/v L-arabinose and incubated at 26°C overnight. The cultures were then pelleted and stored at –20°C. On the day of purification, the pellets were resuspended in a solution of 50 mM NaH_2_PO_4_ pH 8.0, 300 mM NaCl, 10 mM imidazole, 0.1 mg/ml lysozyme, 1 mM PMSF, and 0.1% v/v Triton X-100. After incubating on ice for 1 hour, the cells were disrupted by sonication and cell debris was cleared by centrifugation. The lysate was then passed through a 0.8 µm filter. The clarified lysate was introduced to Ni-NTA resin, which was washed with 10 column volumes of wash buffer 1 (50 mM NaH_2_PO_4_ pH 8.0, 300 mM NaCl, 20 mM imidazole), followed by 10 column volumes of wash buffer 2 (50 mM NaH_2_PO_4_ pH 8.0, 300 mM NaCl, 20 mM imidazole, 10% v/v glycerol). Protein elution was performed using a stepwise increase of imidazole concentrations (100 mM and 250 mM) over ten column volumes. Fractions were analyzed by SDS-PAGE to assess purity. Fractions containing the Cas10-Csm complex were combined and placed on a 5-20% (w/v) sucrose gradient in a solution of 50 mM Tris HCl pH 8.0, 150 mM NaCl, and 5% (v/v) glycerol. Ultracentrifugation was conducted for 41 hours at 31k rpm using a Beckman SW-32TI rotor. Fractions containing the Cas10-Csm complex were identified by A280. The purity of the samples was evaluated by a 4-20% SDS-PAGE gel. The final elutions were concentrated with a Pall centrifugal device with a 30 kDa MWCO membrane, flash-frozen, and stored at −80°C.

### crRNA extraction and validation

crRNAs were isolated from purified Cas10-Csm variants using extraction two times with phenol-chloroform-isoamyl alcohol (25:24:1) and one time with chloroform. The final aqueous layer was collected and a solution of 70% ice-cold ethanol and 300 mM of sodium acetate pH 5.2 was added. The sample was then incubated at –20°C overnight. The next day, the sample was centrifuged, and the supernatant was disposed of. The resulting pellet was rinsed with ethanol and dried down. The RNA was resuspended in water, and its concentration was measured using a Nanodrop spectrometer.

To visualize the crRNA, the extracted RNA was radiolabeled using γ-32P-ATP, followed by separation on a 12% acrylamide urea-PAGE gel. The gels were exposed to a storage phosphor screen and imaged with a Typhoon FLA 7000 imager.

### Saturation affinity binding assays

RNA binding affinity was assessed using fluorescence polarization (FP). Four 43-nucleotide RNAs (Cognate-F, non-complementary-F, MM1-F, and MM2-F) based on the *Nes* target sequence, labeled with 5’ fluorescein and purified by PAGE were obtained from Horizon Discovery. Serial dilutions of Cas10-Csm with final concentrations from 1000-0.48 nM were combined with the fluorescein-labeled RNA and incubated for 2 hours at room temperature before FP was measured on a Bio-Tek Synergy H1 plate reader. The assay was performed in a buffer containing 50 mM Tris-HCl pH 8.0, 400 mM NH_4_Cl, 5% v/v glycerol, 0.025% w/v BSA, and 1 mM EDTA. Two replicate titrations were performed for each target RNA except the non-complementary negative control. No binding was evident for the 0.48 nM and 0.97 nM titration points, so the mean of these FP values was used to baseline correct the data. Data were fit by non-linear regression in GraphPad Prism 9.0 using the expression y= y_max × {R + x + Kd− [(R + x + Kd)^2^− 4Rx]^1/2^}/2R. The fluorescein target RNA is R = 10 nM. Y_max was estimated from a fit of the 1000-62 nM titration points to a hyperbola.

### Target mismatch library

*In vitro* transcription was done using the Promega RiboMax kit with DNA templates obtained from Eurofins. DNA templates for cognate and +1 to +11 mismatches (excluding +6 mismatch) were annealed and extended using Q5 DNA polymerase to create double stranded DNA. For a +6 mismatch, two strands of DNA were combined, heated at 95°C for 5 min and then slowly cooled to room temperature to allow for annealing. Cleanup of the transcription products involved DNase I digestion, phenol-chloroform extraction with phenol-chloroform-isoamyl alcohol 25:24:1, and ethanol precipitation according to the manufacturer’s instructions. The final product was passed through a G-25 column to remove residual NTPs. The quality of the purified RNAs was verified by a 12% acrylamide urea-PAGE run at 150 V for 45 min, stained with SYBR green II, and imaged on a Typhoon FLA 7000 imager.

### COA synthesis assays

Three replicate reactions were performed in all cases. Reactions were performed with 100 nM Cas10-Csm and 400 nM target RNA in TNG buffer (50 mM Tris-HCl pH 8.0, 150 mM NH_4_Cl, 5% v/v glycerol), 500 μM ATP, 30 nM α-^32^P-ATP 3000 Ci/mmol, and 10 mM MgCl_2_ at 37°C for 1 hour. COA products were separated by thin-layer chromatography (TLC) using previously described methods (Rouillon et al. 2019). Briefly, a chamber containing TLC running buffer, composed of 0.2 M ammonium bicarbonate pH 9.3, 70% ethanol, and 30% water was pre-warmed to 35°C to saturate the chamber with buffer vapor. Samples were spotted on a silica TLC plate 1.5 cm from the bottom of the plate and allowed to air-dry. The plate was placed in the chamber for 2 hrs, or until the solvent front was approximately 5 cm from the top. The plate was removed from the chamber, dried with a gentle air stream and cOAs were visualized by phosphorimaging on a Typhoon FLA 7000 imager. The two prominent cOA products were quantified by densitometry using ImageQuant software. A standard curve of serial dilutions of α-^32^P-ATP was also run on the TLC plate and subjected to phosphorimaging. The standard curve was used to ensure our cOA products were within the linear range of the equipment and the slope extracted from linear regression of the curve was used to convert densitometry values to μM cOA (Figure S3). Log_2_ fold change was calculated by taking log_2_ of the relative amount of cOA synthesized in the presence of a mismatched target divided by the amount synthesized with the cognate target (*e.g.* log_2_ [cOA_K524E, +2_ / cOA_K524E, cognate_]). P-values were calculated with a two-tailed t-test for two samples of unequal variance. Uncropped images of the TLC plates are given Figures S4 and S5.

## Acknowledgements

Research reported in this publication was supported by the National Institute of General Medical Sciences of the National Institutes of Health under Award Number R35GM142966 to J.A.D.

