## Supplemental material for "Cas10 residues lining the target RNA binding channel regulate interference by distinguishing cognate target RNA from mismatched targets"

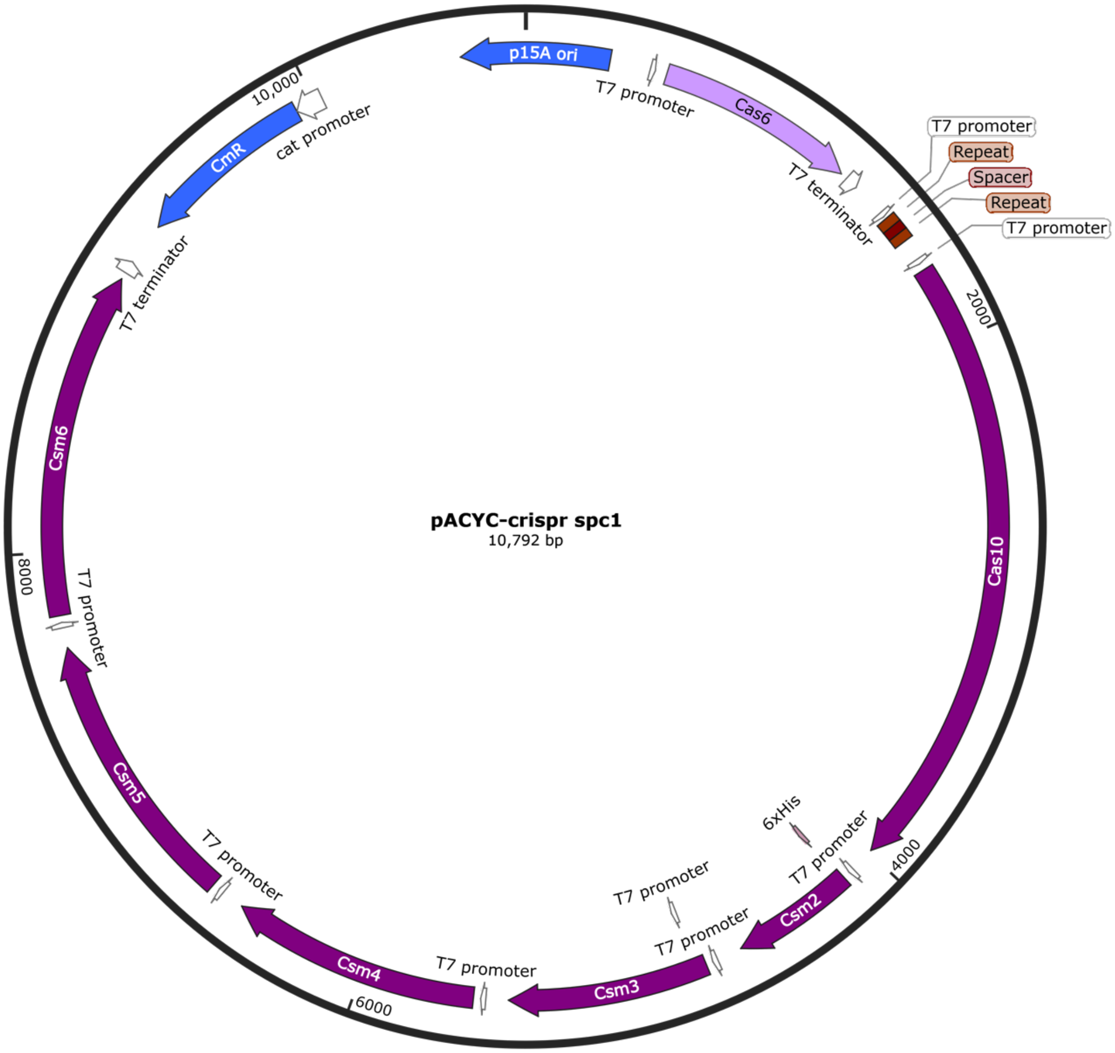


**Figure S1. The map pACYC-CRISPR-spc1.**

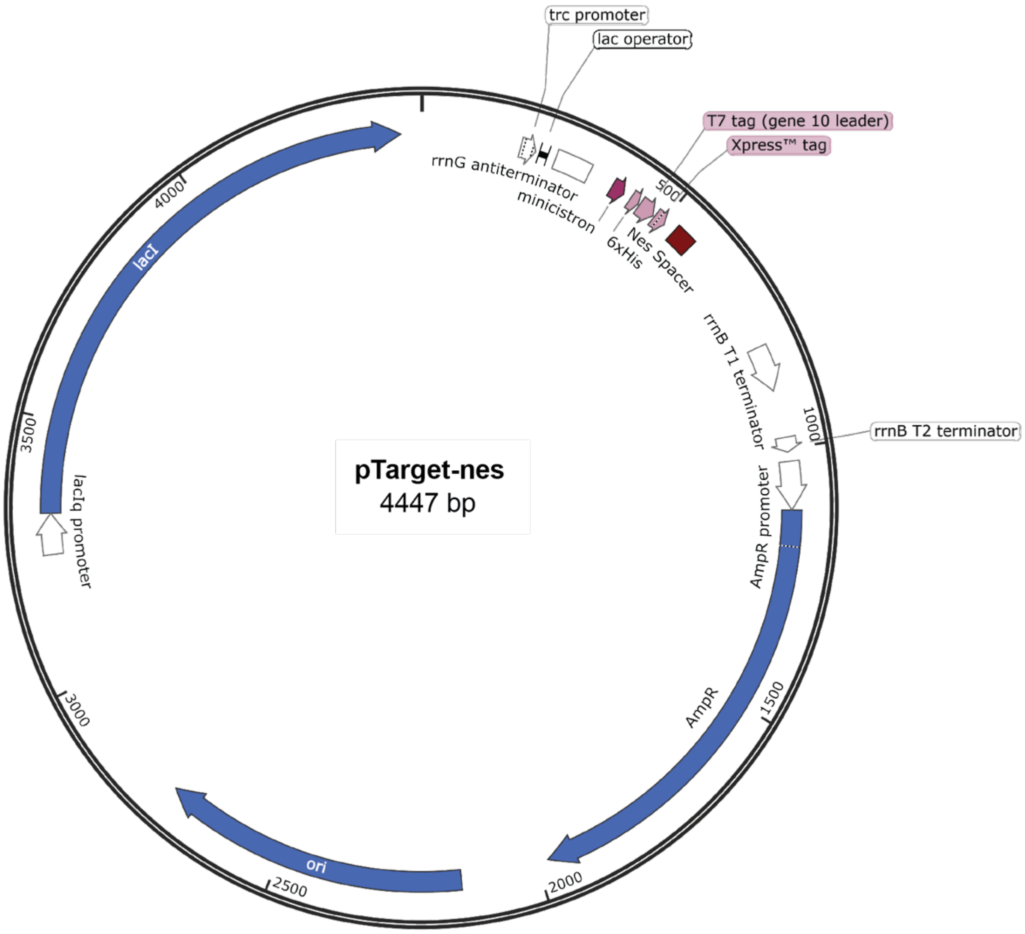


**Figure S2. The map of pTarget-nes.**

**Table S1. Plasmid sequences, gene fragments and oligos used in the study.**

| Name | Sequence | Description |
|---|---|---|
| pr_K524E_F | GCGTCTGGGTGTTGTTCGTGCA | Fwd primer to introduce K524E mutation into pACYC-CRISPR-spc1 |
| pr_K524E_R | TCAACACCTTCTTCACGATCAACATAG | Rev primer to introduce K524E mutation into pACYC-CRISPR-spc1 |
| pr_K628E_F | GTATCCGGTTAGCAAAATGGCC | Fwd primer to introduce K628E mutation into pACYC-CRISPR-spc1 |
| pr_K628E_R | TCACCGCTAAACATGCCAACACC | Rev primer to introduce K628E mutation into pACYC-CRISPR-spc1 |
| pr_K691E_F | GGCCTTCATTTACAAAATGCTG | Fwd primer to introduce K691E mutation into pACYC-CRISPR-spc1 |
| pr_K691E_R | TCACCATGTTCATCGGTCTGGCT | Rev primer to introduce K691E mutation into pACYC-CRISPR-spc1 |
| pr_MAB_017 | AGCCTTCATTGAAAAAATGCTGG | Fwd primer to introduce Y695E mutation into pACYC-CRISPR-spc1 |
| pr_MAB_018 | TTACCATGTTCATCGG | Rev primer to introduce Y695E mutation into pACYC-CRISPR-spc1 |
| pr_K691Y695E_F | CATTGAGAAAATGCTGGCACTGCTG | Fwd primer to introduce K691E and Y695E mutations into pACYC-CRISPR-spc1 |
| pr_K691Y695E_R | AAGGCCTCACCATGTTCATCGGTCTG | Rev primer to introduce K691E and Y695E mutations into pACYC-CRISPR-spc1 |
| pr_R754E_F | GGAAGCCGACTAACATATGGCT | Fwd primer to introduce R754E mutation into pACYC-CRISPR-spc1 |
| pr_R754E_R | TCGATCTGATAGATATAGTATTCCAGGG | Rev primer to introduce R754E mutation into pACYC-CRISPR-spc1 |
| pr_SAK_011 | GCCTGATACAGATTAAATCAGAAC | Primer to linearize pTRC to excise spacer coding region. GA |
| pr_SAK_012 | GTCATCGTCGTACAGATCC | Primer to linearize pTRC to excise spacer coding region. GA |
| pr_SAK_047 | ATGCAAATGAACCGGCTTACTACGAAGGCGGC | Fwd primer to introduce mismatch of segment 2 on pTarget-cn20 with Q5 PCR MMS2 |
| pr_SAK_048 | ATGTTACATCGATAAGCTTG | Rev primer to introduce mismatch of segment 2 on pTarget-cn20 with Q5 PCR MMS2 |

| | | |
|---|---|---|
| pACYC_Cas10_5x | CAGAATAATAACCAGGTTCGCATCTATAGCAAAAACAAACCGTATATTGGCATTGGCATTAGCACCAATCTGTGGATGTGTGATTATGATTATGCAAGCCAGAATCAGGATATGCGCGAAAAAGGTATTGGTAGCTATGTTGATCGTGAAGAAGGTGTTGAGCGTCTGGGTGTTGTTCGTGCAGATATTGATAATCTGGGTGCAACCTTTATTAGCGGCATTCCGGAAAAATACAATAGCATTAGCCGTACCGCAACCCTGAGCCGTCAGCTGAGTCTGTTCTTTAAATACGAGCTGAACCATCTGCTGGAAAACTATCAGATTACCGCAATTTATAGTGGCGGAGATGACCTGTTTCTGATTGGTGCATGGGATGATATTATCGAAGCGAGCATTTACATCAACGATAAATTCAAAGAGTTTACCCTGGACAAACTGACCCTGAGTGCCGGTGTTGGCATGTTTAGCGGTGAGTATCCGGTTAGCAAAATGGCCTTTGAGACAGGTCGTCTGGAAGAGGCAGCAAAAACTGGCGAAAAAAACCAGATTAGTCTGTGGCTGCAAGAGAAAGTGTATAACTGGGATGAGTTCAAAAAAAACATTCTGGAAGAGAAACTGCTGGTTCTGCAGCAGGGTTTTAGCCAGACCGATGAACATGGTGAGGCCTTCATTGAGAAAATGCTGGCACTGCTGCGTAATAACGAAGCAATTAACATTGCACGTCTGGCATACCTGCTGGCACGTAGTAAAATGAATGAAGATTTCACCAGCAAAATCTTTAACTGGGCACAGAACGACAAAGACAAAAATCAACTGATTACAGCCCTGGAATACTATATCTATCAGATCGAAGAAGCCGACTAACATATGGCTGCGTGG | Gene segment to introduce 5x mutation into pACYC-CRISPR-spc1 |
| pTRC_nes_MMS2 | GGGATCTGTACGACGATGACGATAAGGATCCAACCCTTTTCCAAGCTTCTTTGTACTGATGATTTATATACAAGCCGATACGTCAAGAGCAGCATGCTTCCAAGGCGAATTCGAAGCTTGGCTGTTTTGGCGGATGAGAGAAGATTTTCAGCCTGATACAGATTAAATCA | Gene segment to clone MMS2 from +7 to +12 into pTarget-nes |
| CN20 pACYC gene segment | CACTATAGGGAGACCATGGGATCGATACCCACCCCGAAGAAAAGGGGACGAGAACTAGTAATAATTGTCATTTGCATACGTTACATCGATGATCGATACCCACCCCGAAGAAAAGGGGACGAGAACCTCGAGGCTGTGGTCTAGACATTC | Cn20 ThermoFisher GeneArt |
| prJAD_500A | GAATTCTAATACGACTCACTATAGGCTTTGTACTGATGATTTATATACTTCGGCA | DNA forward template for Q5 PCR extension to introduce a mismatch from cog, +1 to +5 |
| prJAD_500B | TTTAGAGAACGTATGCCGAAGTATATAAATC | DNA reverse template for Q5 PCR extension for cognate |
| prJAD_501 | TTTAGAGATCGTATGCCGAAGTATATAAATC | DNA reverse template for Q5 PCR extension to introduce a mismatch at the +1 position of nes |
| prJAD_502 | TTTAGAGAAGGTATGCCGAAGTATATAAATC | DNA reverse template for Q5 PCR extension to introduce a mismatch at the +2 position of nes |

| | | |
|---|---|---|
| prJAD_503 | TTTAGAGAACCTATGCCGAAGTATATAAATC | DNA reverse template for Q5 PCR extension to introduce a mismatch at the +3 position of nes |
| prJAD_504 | TTTAGAGAACGAATGCCGAAGTATATAAATC | DNA reverse template for Q5 PCR extension to introduce a mismatch at the +4 position of nes |
| prJAD_505 | TTTAGAGAACGTTTGCCGAAGTATATAAATC | DNA reverse template for Q5 PCR extension to introduce a mismatch at the +5 position of nes |
| prMON_906S | GAATTCTAATACGACTCACTATAGGCTTTGTACTG ATGATTTATATACTTCGGCTTACGTTCTCTAAA | DNA sense strand template to introduce a +6 mismatch along the nes target RNA |
| prMON_906AS | TTTAGAGAACGTAAGCCGAAGTATATAAATCATCA GTACAAAGCCTATAGTGAGTCGTATTAGAATTC | DNA antisense strand template to introduce a +6 mismatch along the nes target RNA |
| prMON_507A | GAATTCTAATACGACTCACTATAGGCTTTGTACTG ATGATTTATATACTTCGGGA | DNA forward template for Q5 PCR extension to introduce a mismatch at the +7 position of nes |
| prMON_507B | TTTAGAGAACGTATCCCGAAGTATATAAATC | DNA reverse template for Q5 PCR extension to introduce a mismatch at the +7 position of nes |
| prMON_508A | GAATTCTAATACGACTCACTATAGGCTTTGTACTG ATGATTTATATACTTCGCCA | DNA forward template for Q5 PCR extension to introduce a mismatch at the +8 position of nes |
| prMON_508B | TTTAGAGAACGTATGGCGAAGTATATAAATC | DNA reverse template for Q5 PCR extension to introduce a mismatch at the +8 position of nes |
| prMON_509A | GAATTCTAATACGACTCACTATAGGCTTTGTACTG ATGATTTATATACTTCCGCA | DNA forward template for Q5 PCR extension to introduce a mismatch at the +9 position of nes |
| prMON_509B | TTTAGAGAACGTATGCGGAAGTATATAAATC | DNA reverse template for Q5 PCR extension to introduce a mismatch at the +9 position of nes |
| prMON_510A | GAATTCTAATACGACTCACTATAGGCTTTGTACTG ATGATTTATATACTTGGGCA | DNA forward template for Q5 PCR extension |

| | | |
|---|---|---|
| | | to introduce a mismatch at the +10 position of nes |
| prMON_510B | TTTAGAGAACGTATGCCCAAGTATATAAATC | DNA reverse template for Q5 PCR extension to introduce a mismatch at the +10 position of nes |
| prMON_511A | GAATTCTAATACGACTCACTATAGGCTTTGTACTG<br>ATGATTTATATACTACGGCA | DNA forward template for Q5 PCR extension to introduce a mismatch at the +11 position of nes |
| prMON_511B | TTTAGAGAACGTATGCCGTAGTATATAAATC | DNA reverse template for Q5 PCR extension to introduce a mismatch at the +11 position of nes |
| pmRNA01-F | CUUUGUACUGAUGAUUUAUAUACUUCGGCAUAC<br>GUUCUCUAAA | Nes target sequence, labeled with 5' fluorescein |
| pmRNA01-FMM1 | CUUUGUACUGAUGAUUUAUAUACUUCGGCUAUG<br>CAUCUCUAAA | Nes target sequence, with nucleotides +1 to +6 converted to their corresponding Watson-Crick pairs, labeled with a 5' fluorescein |
| pmRNA01-FMM2 | CUUUGUACUGAUGAUUUAUAUACAAGCCGAUAC<br>GUUCUCUAAA | Nes target sequence, with nucleotides +7 to +12 converted to their corresponding Watson-Crick pairs, labeled with a 5' fluorescein |
| pmRNA-FNC | GCUGACAUUAAGAUUACUAUUUAUUAUCCUUGC<br>GAUUUCUACG | Non-complementary RNA target sequence, labeled with a 5' fluorescein |
| >pTarget-nes (4447 bp)<br>GTTTGACAGCTTATCATCGACTGCACGGTGCACCAATGCTTCTGGCGTCAGGCAGCCATCGGAAG<br>CTGTGGTATGGCTGTGCAGGTCGTAAATCACTGCATAATTCGTGTCGCTCAAGGCGCACTCCCGT<br>TCTGGATAATGTTTTTTGCGCCGACATCATAACGGTTCTGGCAAATATTCTGAAATGAGCTGTTGA<br>CAATTAATCATCCGGCTCGTATAATGTGTGGAATTGTGAGCGGATAACAATTTCACACAGGAAACA<br>GCGCCGCTGAGAAAAAGCGAAGCGGCACTGCTCTTTAACAATTTATCAGACAATCTGTGTGGGCA<br>CTCGACCGGAATTATCGATTAACTTTATTATTAAAAATTAAAGAGGTATATATTAATGTATCGATTAA<br>ATAAGGAGGAATAAACCATGGGGGGTTCTCATCATCATCATCATCATGGTATGGCTAGCATGACTG<br>GTGGACAGCAAATGGGTCGGGATCTGTACGACGATGACGATAAGGATCCAACCCTTTTCCAAGCT<br>TCTTTGTACTGATGATTTATATACTTCGGCATACGTCAAGAGCAGCATGCTTCCAAGGCGAATTCG<br>AAGCTTGGCTGTTTTGGCGGATGAGAGAAGATTTTCAGCCTGATACAGATTAAATCAGAACGCAGA<br>AGCGGTCTGATAAAACAGAATTTGCCTGGCGGCAGTAGCGCGGTGGTCCCACCTGACCCCATGC<br>CGAACTCAGAAGTGAAACGCCGTAGCGCCGATGGTAGTGTGGGGTCTCCCCATGCGAGAGTAGG<br>GAACTGCCAGGCATCAAATAAAACGAAAGGCTCAGTCGAAAGACTGGGCCTTTCGTTTTATCTGTT<br>GTTTGTCGGTGAACGCTCTCCTGAGTAGGACAAATCCGCCGGGAGCGGATTTGAACGTTGCGAA<br>GCAACGGCCCGGAGGGTGGCGGGCAGGACGCCCGCCATAAACTGCCAGGCATCAAATTAAGCA<br>GAAGGCCATCCTGACGGATGGCCTTTTTGCGTTTCTACAAACTCTTTTGTTTATTTTTCTAAATACA<br>TTCAAATATGTATCCGCTCATGAGACAATAACCCTGATAAATGCTTCAATAATATTGAAAAAGGAAG | | |

AGTATGAGTATTCAACATTTCCGTGTCGCCCTTATTCCCTTTTTTGCGGCATTTTGCCTTCCTGTTT
TTGCTCACCCAGAAACGCTGGTGAAAGTAAAAGATGCTGAAGATCAGTTGGGTGCACGAGTGGGT
TACATCGAACTGGATCTCAACAGCGGTAAGATCCTTGAGAGTTTTCGCCCCGAAGAACGTTTTCCA
ATGATGAGCACTTTTAAAGTTCTGCTATGTGGCGCGGTATTATCCCGTGTTGACGCCGGGCAAGA
GCAACTCGGTCGCCGCATACACTATTCTCAGAATGACTTGGTTGAGTACTCACCAGTCACAGAAAA
GCATCTTACGGATGGCATGACAGTAAGAGAATTATGCAGTGCTGCCATAACCATGAGTGATAACAC
TGCGGCCAACTTACTTCTGACAACGATCGGAGGACCGAAGGAGCTAACCGCTTTTTTGCACAACA
TGGGGGATCATGTAACTCGCCTTGATCGTTGGGAACCGGAGCTGAATGAAGCCATACCAAACGAC
GAGCGTGACACCACGATGCCTGTAGCAATGGCAACAACGTTGCGCAAACTATTAACTGGCGAACT
ACTTACTCTAGCTTCCCGGCAACAATTAATAGACTGGATGGAGGCGGATAAAGTTGCAGGACCAC
TTCTGCGCTCGGCCCTTCCGGCTGGCTGGTTTATTGCTGATAAATCTGGAGCCGGTGAGCGTGG
GTCTCGCGGTATCATTGCAGCACTGGGGCCAGATGGTAAGCCCTCCCGTATCGTAGTTATCTACA
CGACGGGGAGTCAGGCAACTATGGATGAACGAAATAGACAGATCGCTGAGATAGGTGCCTCACT
GATTAAGCATTGGTAACTGTCAGACCAAGTTTACTCATATATACTTTAGATTGATTTAAAACTTCATT
TTTAATTTAAAAGGATCTAGGTGAAGATCCTTTTTGATAATCTCATGACCAAAATCCCTTAACGTGA
GTTTTCGTTCCACTGAGCGTCAGACCCCGTAGAAAAGATCAAAGGATCTTCTTGAGATCCTTTTTTT
CTGCGCGTAATCTGCTGCTTGCAAACAAAAAAACCACCGCTACCAGCGGTGGTTTGTTTGCCGGA
TCAAGAGCTACCAACTCTTTTTCCGAAGGTAACTGGCTTCAGCAGAGCGCAGATACCAAATACTGT
CCTTCTAGTGTAGCCGTAGTTAGGCCACCACTTCAAGAACTCTGTAGCACCGCCTACATACCTCG
CTCTGCTAATCCTGTTACCAGTGGCTGCTGCCAGTGGCGATAAGTCGTGTCTTACCGGGTTGGAC
TCAAGACGATAGTTACCGGATAAGGCGCAGCGGTCGGGCTGAACGGGGGGTTCGTGCACACAGC
CCAGCTTGGAGCGAACGACCTACACCGAACTGAGATACCTACAGCGTGAGCTATGAGAAAGCGC
CACGCTTCCCGAAGGGAGAAAGGCGGACAGGTATCCGGTAAGCGGCAGGGTCGGAACAGGAGA
GCGCACGAGGGAGCTTCCAGGGGGAAACGCCTGGTATCTTTATAGTCCTGTCGGGTTTCGCCAC
CTCTGACTTGAGCGTCGATTTTTGTGATGCTCGTCAGGGGGGCGGAGCCTATGGAAAAACGCCAG
CAACGCGGCCTTTTTACGGTTCCTGGCCTTTTGCTGGCCTTTTGCTCACATGTTCTTTCCTGCGTT
ATCCCCTGATTCTGTGGATAACCGTATTACCGCCTTTGAGTGAGCTGATACCGCTCGCCGCAGCC
GAACGACCGAGCGCAGCGAGTCAGTGAGCGAGGAAGCGGAAGAGCGCCTGATGCGGTATTTTCT
CCTTACGCATCTGTGCGGTATTTCACACCGCATATGGTGCACTCTCAGTACAATCTGCTCTGATGC
CGCATAGTTAAGCCAGTATACACTCCGCTATCGCTACGTGACTGGGTCATGGCTGCGCCCCGACA
CCCGCCAACACCCGCTGACGCGCCCTGACGGGCTTGTCTGCTCCCGGCATCCGCTTACAGACAA
GCTGTGACCGTCTCCGGGAGCTGCATGTGTCAGAGGTTTTCACCGTCATCACCGAAACGCGCGA
GGCAGCAGATCAATTCGCGCGCGAAGGCGAAGCGGCATGCATTTACGTTGACACCATCGAATGG
TGCAAAACCTTTCGCGGTATGGCATGATAGCGCCCGGAAGAGAGTCAATTCAGGGTGGTGAATGT
GAAACCAGTAACGTTATACGATGTCGCAGAGTATGCCGGTGTCTCTTATCAGACCGTTTCCCGCG
TGGTGAACCAGGCCAGCCACGTTTCTGCGAAAACGCGGGAAAAAGTGGAAGCGGCGATGGCGG
AGCTGAATTACATTCCCAACCGCGTGGCACAACAACTGGCGGGCAAACAGTCGTTGCTGATTGGC
GTTGCCACCTCCAGTCTGGCCCTGCACGCGCCGTCGCAAATTGTCGCGGCGATTAAATCTCGCG
CCGATCAACTGGGTGCCAGCGTGGTGGTGTCGATGGTAGAACGAAGCGGCGTCGAAGCCTGTAA
AGCGGCGGTGCACAATCTTCTCGCGCAACGCGTCAGTGGGCTGATCATTAACTATCCGCTGGATG
ACCAGGATGCCATTGCTGTGGAAGCTGCCTGCACTAATGTTCCGGCGTTATTTCTTGATGTCTCTG
ACCAGACACCCATCAACAGTATTATTTTCTCCCATGAAGACGGTACGCGACTGGGCGTGGAGCAT
CTGGTCGCATTGGGTCACCAGCAAATCGCGCTGTTAGCGGGCCCATTAAGTTCTGTCTCGGCGC
GTCTGCGTCTGGCTGGCTGGCATAAATATCTCACTCGCAATCAAATTCAGCCGATAGCGGAACGG
GAAGGCGACTGGAGTGCCATGTCCGGTTTTCAACAAACCATGCAAATGCTGAATGAGGGCATCGT
TCCCACTGCGATGCTGGTTGCCAACGATCAGATGGCGCTGGGCGCAATGCGCGCCATTACCGAG
TCCGGGCTGCGCGTTGGTGCGGATATCTCGGTAGTGGGATACGACGATACCGAAGACAGCTCAT
GTTATATCCCGCCGTTAACCACCATCAAACAGGATTTTCGCCTGCTGGGGCAAACCAGCGTGGAC
CGCTTGCTGCAACTCTCTCAGGGCCAGGCGGTGAAGGGCAATCAGCTGTTGCCCGTCTCACTGG
TGAAAAGAAAAACCACCCTGGCGCCCAATACGCAAACCGCCTCTCCCCGCGCGTTGGCCGATTCA
TTAATGCAGCTGGCACGACAGGTTTCCCGACTGGAAAGCGGGCAGTGAGCGCAACGCAATTAAT
GTAAGTTAGCGCGAATTGATCTG

>pACYC-CRISPR-spc1 (10792 bp)
ATGCACGAACCCCCCGTTCAGTCCGACCGCTGCGCCTTATCCGGTAACTATCGTCTTGAGTCCAA
CCCGGAAAGACATGCAAAAGCACCACTGGCAGCAGCCACTGGTAATTGATTTAGAGGAGTTAGTC

TTGAAGTCATGCGCCGGTTAAGGCTAAACTGAAAGGACAAGTTTTGGTGACTGCGCTCCTCCAAG
CCAGTTACCTCGGTTCAAAGAGTTGGTAGCTCAGAGAACCTTCGAAAAACCGCCCTGCAAGGCGG
TTTTTTCGTTTTCAGAGCAAGAGATTACGCGCAGACCAAAACGATCTCAAGAAGATCATCTTATTAA
TCAGATAAAATATTTCTAGATTTCAGTGCAATTTATCTCTTCAAATGTAGCACCTGAAGTCAGCCCC
ATACGATATAAGTTGTAATTCTCATGTTTGACAGCCTTCAGATCCGATAGACTAGCCGCTGGTAATA
ATACGACTCACTATAGGGAGAGAATTCTAAGACCGAAAGTCGGAAACAAAGAGGATTTATATGATC
AACAAAATCACCGTGGAACTGGATCTGCCGGAAAGCATTCGTTTTCAGTATCTGGGTAGCGTTCTG
CATGGTGTTCTGATGGATTATCTGAGTGATGATATTGCAGATCAGCTGCATCACGAATTTGCATAT
AGTCCGCTGAAACAGCGCATCTACCACAAAAACAAAAAAATCATCTGGGAAATCGTGTGCATGAG
CGATAACCTGTTTAAAGAAGTGGTGAAACTGTTTAGCAGCAAAAATAGCCTGCTGCTGAAATATTA
CCAGACCAACATTGATATCCAGAGCTTCCAGATCGAAAAAATCAATGTGCAGAACATGATGAATCA
GCTGCTGCAGGTTGAGGATCTGAGCCGTTATGTTCGTCTGAACATTCAGACCCCGATGAGCTTCA
AATATCAGAACAGCTATATGATCTTCCCGGATGTGAAACGTTTTTTCCGCAGCATTATGATTCAGTT
CGATGCCTTTTTTGAAGAATACCGCATGTACGATAAAGAAACCCTGAACTTCCTGGAAAAAAACGT
GAACATCGTGGATTATAAACTGAAAAGCACCCGCTTTAATCTGGAAAAAGTTAAAATTCCGAGCTTT
ACCGGTGAGATCGTGTTCAAAATCAAAGGTCCGCTGCCGTTTCTGCAGCTGACCCATTTTCTGCT
GAAATTTGGTGAATTTAGCGGCAGCGGTATTAAAACCAGCCTGGGTATGGGTAAATATAGCATCAT
CTAAAAGCTTTCCTGTGAGCAGCGAAAGCCTAGCATAACCCCTTGGGGCCTCTAAACGGGTCTTG
AGGGGTTTTTTGTTATACGCGAGATAATCACTTGCATAGCTGCGTATGGAGGAAGCAACTCTTGAG
TGTTAATATGTTGACCCCTGTATTAGGGATGCGGGTAGTAGATGTGGGCAGAGACACCCACACTG
CCAGATCTTAATACGACTCACTATAGGGAGACCATGGGATCGATACCCACCCCGAAGAAAAGGGG
ACGAGAACACGTATGCCGAAGTATATAAATCATCAGTACAAAGGATCGATACCCACCCCGAAGAA
AAGGGGACGAGAACCTCGAGGCTGTGGTCTAGACATTCCATACATATCGGGGGGGTAGGGGTTT
TTTGTGTGCCTCTAGTGGCTGGCTAAGAATAATACGACTCACTATAGGGAGAGGATCCATAAAGGA
GGTAAATAATGAACAAAAAAAACATCCTGATGTATGGCAGCCTGCTGCATGATATTGGCAAAATTA
TCTATCGTAGCGGTGATCATACCTTTAGCCGTGGCACCCATAGCAAACTGGGTCATCAGTTTCTGA
GCCAGTTTAGCGAATTTAAAGATAACGAAGTGCTGGATAACGTGGCCTATCATCATTATAAAGAAC
TGGCAAAAGCCAACCTGGATAATGATAATACCGCCTACATTACCTATATCGCCGATAATATTGCAA
GCGGTATTGATCGTCGCGATATTATTGAAGAGGGTGATGAAGAATATGAGAAACAACTGTTCAACT
TCGATAAATACACACCGCTGTATAGCGTGTTTAACATTGTGAATAGCGAAAAACTGAAACAGACCA
ACGGCAAATTCAAATTTAGCAACGAAAGCAACATCGAATACCCGAAAACCGAAAACATTCAGTATA
GCAGCGGTAATTATACCACCCTGATGAAAGATATGAGCCATGATCTGGAACATAAACTGAGCATTA
AAGAAGGCACCTTTCCGAGTCTGCTGCAGTGGACCGAAAGCCTGTGGCAGTATGTTCCGAGCAG
CACCAATAAAAACCAGCTGATTGATATCAGCCTGTATGACCATAGCCGTATTACCTGTGCAATTGC
CAGCTGCATTTTTGATTATCTGAACGAGAACAACATCCACAACTATAAAGATGAACTGTTTAGCAAA
TATGAAAACACCAAATCCTTTTATCAGAAAGAGGCATTTCTGCTGCTGAGCATGGATATGAGCGGT
ATTCAGGATTTCATCTATAACATTAGCGGTAGCAAAGCACTGAAAAGCCTGCGTAGCCGTAGCTTT
TATCTGGAACTGATGCTGGAAGTTATTGTTGATCAGCTGCTGGAACGCCTGGAACTGGCACGTGC
AAATCTGCTGTATACCGGTGGTGGTCATGCATATCTGCTGGTTAGCAATACCGACAAAGTGAAAAA
AAAAATCACCCAGTTCAACAACGAACTGAAAAAATGGTTTATGAGCGAGTTTACCACCGATCTGAG
CCTGTCAATGGCATTTGAAAAATGTAGTGGTGATGACCTGATGAATACCAGCGGCAATTATCGTAC
CATTTGGCGTAATGTTAGCAGCAAACTGAGCGATATTAAAGCCCACAAATATAGCGCAGAGGACAT
TCTGAAACTGAACCATTTTCATAGTTATGGCGATCGCGAATGTAAAGAATGTCTGCGTAGCGATAT
TGACATTAACGATGATGGTCTGTGTAGCATTTGCGAAGGCATTATTAACATCAGCAATGATCTGCG
CGACAAATCGTTTTTTGTGCTGAGCGAAACCGGTAAACTGAAAATGCCGTTTAACAAATTCATCAG
CGTGATCGATTATGAAGAGGCCGAAATGCTGGTTCAGAATAATAACCAGGTTCGCATCTATAGCAA
AAACAAACCGTATATTGGCATTGGCATTAGCACCAATCTGTGGATGTGTGATTATGATTATGCAAG
CCAGAATCAGGATATGCGCGAAAAAGGTATTGGTAGCTATGTTGATCGTGAAGAAGGTGTTAAAC
GTCTGGGTGTTGTTCGTGCAGATATTGATAATCTGGGTGCAACCTTTATTAGCGGCATTCCGGAAA
AATACAATAGCATTAGCCGTACCGCAACCCTGAGCCGTCAGCTGAGTCTGTTCTTTAAATACGAGC
TGAACCATCTGCTGGAAAACTATCAGATTACCGCAATTTATAGTGGCGGAGATGACCTGTTTCTGA
TTGGTGCATGGGATGATATTATCGAAGCGAGCATTTACATCAACGATAAATTCAAAGAGTTTACCC
TGGACAAACTGACCCTGAGTGCCGGTGTTGGCATGTTTAGCGGTAAATATCCGGTTAGCAAAATG
GCCTTTGAGACAGGTCGTCTGGAAGAGGCAGCAAAAACTGGCGAAAAAAACCAGATTAGTCTGTG
GCTGCAAGAGAAAGTGTATAACTGGGATGAGTTCAAAAAAAACATTCTGGAAGAGAAACTGCTGG
TTCTGCAGCAGGGTTTTAGCCAGACCGATGAACATGGTAAAGCCTTCATTTACAAAATGCTGGCAC

TGCTGCGTAATAACGAAGCAATTAACATTGCACGTCTGGCATACCTGCTGGCACGTAGTAAAATGA
ATGAAGATTTCACCAGCAAAATCTTTAACTGGGCACAGAACGACAAAGACAAAAATCAACTGATTA
CAGCCCTGGAATACTATATCTATCAGATCCGTGAAGCCGACTAACATATGGCTGCGTGGTCAAATG
TGCGTACCCTAACCCCTTCCCCGGTCAATCGGGGCGGATGGGGTTTTTTGTGCGTACTTCATTAT
GTATATTAATACGACTCACTATAGGGAGAAGATCTATAAAGGAGGTAAATAATGGGTCACCACCAT
CATCACCATAGCGGTGGAATTCTGGCCAAAACCAAAAGCGGCAAAACCATTGATCTGACCTTTGC
ACATGAAGTGGTTAAAAGCAATGTGAAAAACGTGAAAGACCGCAAAGGCAAAGAAAAACAGGTTC
TGTTTAATGGTCTGACCACCAGTAAACTGCGTAATCTGATGGAACAGGTTAATCGCCTGTATACCA
TTGCCTTTAATAGCAATGAAGATCAGCTGAACGAAGAGTTTATCGATGAACTGGAATATCTGAAAAT
CAAATTCTACTATGAAGCCGGTCGTGAGAAAAGCGTTGATGAGTTTCTGAAAAAAACCCTGATGTT
CCCGATTATTGATCGCGTGATCAAAAAAGAAAGCAAAAAATTCTTCCTGGACTACTGCAAATATTTC
GAAGCACTGGTTGCATACGCCAAATATTACCAGAAAGAGGACTAAACGCGTGCTGCGTGGTCAAA
TGTGCGTAGACCAACCCCTTGCGGCCTCAATCGGGGGGGATGGGGTTTTTTGTCAGGCAAGTCT
CAGCTGGTTTAATACGACTCACTATAGGGAGAGAATTCATAAAGGAGGTAAATAATGGGGTTTTTT
GTCAGGCAAGTCTCAGCTGGTTTAATACGACTCACTATAGGGAGAGAATTCCCCAGCAGTATAACA
GGAGGACACCAGATGTACAGCAAAATCAAAATCAGCGGCACCATTGAAGTTGTTACCGGTCTGCA
TATTGGTGGTGGTGGCGAAAGCAGCATGATTGGTGCAATTGATAGTCCGGTTGTTCGTGATCTGC
AGACCAAACTGCCGATTATTCCGGGTAGCAGCATTAAAGGTAAAATGCGTAATCTGCTGGCCAAA
CACTTTGGCCTGAAAATGAAACAAGAAAGCCATAACCAGGATGATGAACGTGTTCTGCGTCTGTTT
GGTAGCAGCGAAAAAGGTAATATTCAGCGTGCTCGCCTGCAGATTAGTGATGCATTTTTTAGCGAA
AAAACCAAAGAACACTTCGCCCAGAATGATATTGCATACACCGAAACCAAATTCGAGAATACCATT
AATCGTCTGACCGCAGTTGCAAATCCGCGTCAGATTGAACGTGTGACCCGTGGTAGCGAATTTGA
CTTTGTGTTTATCTATAACGTGGATGAAGAGTCCCAGGTGGAAGATGATTTTGAAAACATTGAGAA
AGCGATCCATCTGCTGGAAAATGATTATCTGGGTGGCGGTGGTACACGTGGTAATGGTCGTATTC
AGTTTAAAGACACCAACATTGAAACCGTGGTGGGTGAATATGATAGCACCAATCTGAAAATCAAAT
AAAAGCTTACCTGGAGATCAAGGAGATTACTCTAACCCCATCGGCCGTCTTAGGGGTTTTTTGTCC
TGTGTTAGCTGGAGGGTATAATACGACTCACTATAGGGAGACCCGGGATAAAGGAGGTAAATAAT
GACCCTGGCAACCAAAGTTTTTAAACTGAGCTTTAAAACACCGGTGCATTTCGGTAAAAAACGTCT
GAGTGATGGTGAAATGACCATTACCGCAGATACCCTGTTTAGCGCACTGTTTATTGAAACCCTGCA
GCTGGGTAAAGATACCGATTGGCTGCTGAATGATCTGATTATTAGCGATACCTTTCCGTATGAGAA
CGAGCTGTATTATCTGCCGAAACCGCTGATTAAAATCGACAGCAAAGAAGAGGATAACCACAAAG
CCTTCAAAAAACTGAAATATGTGCCGGTGCATCACTATAACCAGTATCTGAATGGTGAACTGAGCG
CAGAAGATGCAACCGATCTGAATGATATTTTCAACATCGGCTATTTCAGCCTGCAGACCAAAGTTA
GCCTGATTGCACAAGAAACCGATAGCAGCGCAGATAGCGAACCGTATAGCGTTGGCACCTTTACC
TTTGAACCGGAAGCAGGTCTGTATTTTATCGCAAAAGGTAGCGAAGAAACCCTGGATCATCTGAAT
AACATTATGACCGCACTGCAGTATAGCGGTCTGGGTGGTAAACGTAATGCAGGTTATGGTCAGTTT
GAGTACGAAATCATTAATAACCAGCAGCTGAGCAAACTGCTGAATCAGAATGGTAAACATAGCATT
CTGCTGAGCACCGCAATGGCAAAAAAAGAAGAAATTGAAAGCGCACTGAAAGAGGCACGTTATAT
TCTGACCAAACGTAGCGGTTTTGTTCAGAGCACCAATTATAGCGAAATGCTGGTGAAAAAAAGCGA
CTTCTATAGCTTTAGCAGCGGCAGCGTTTTCAAAAACATTTTTAACGGCGATATCTTCAACGTGGG
CCATAATGGCAAACATCCGGTTTATCGTTATGCTAAACCGCTGTGGCTGGAAGTTTAATCATGATT
TCTTGTCGAACTGGACAGTAGCAGAACCGCTAACGGGGGCGAAGGGGTTTTTTGTGACATACGAG
CTGATTGAACTAATACGACTCACTATAGGGAGAGGTACCATAAAGGAGGTAAATAATGACCATCAA
AAACTATGAGGTGGTGATTAAAACCCTGGGTCCGATTCATATTGGTAGCGGTCAGGTTATGAAAAA
ACAGGATTATATCTACGACTTTTATAACAGCAAAGTGTATATGATCAACGGCAACAAACTGGTGAA
ATTTCTGAAACGCAAAAACCTGCTGTATACCTATCAGAACTTTCTGCGTTATCCGCCTAAAAATCCG
CGTGAAAATGGTCTGAAAGATTATCTGGATGCCCAGAATGTTAAACAGAGCGAATGGGAAGCATTT
GTGAGCTATAGCGAAAAAGTGAACCAGGGCAAAAAATACGGTAATACCCGTCCGAAACCGCTGAA
TGATCTGCATCTGATGGTTCGTGATGGTCAGAATAAAGTTTATCTGCCTGGTAGCAGCATTAAAGG
TGCAATTAAAACCACCCTGGTGAGCAAATATAACAACGAAAAAAACAAAGATATCTATAGCAAAATC
AAAGTGAGCGATAGCAAACCGATTGATGAAAGCAATCTGGCCATCTATCAGAAAATCGACATCAAC
AAAAGCGAGAAAAGCATGCCGCTGTATCGTGAATGTATTGATGTGAACACCGAGATCAAATTCAAA
CTGACCATCGAGGATGAAATCTACAGCATCAATGAAATCGAACAGAGCATCCAGGACTTCTATAAA
AACTACTATGATAAATGGCTGGTCGGCTTTAAAGAAACCAAAGGTGGTCGTCGTTTTGCACTGGAA
GGTGGTATTCCGGATGTTCTGAATCAAAACATTCTGTTTCTGGGTGCAGGCACCGGTTTTGTGAGC
AAAACCACACATTATCAGCTGAAAAATCGCAAACAGGCCAAACAGGATAGCTTTGAAATTCTGACG

AAAAAATTCCGTGGCACCTACGGCAAAATGAAAGAAATTCCGAGCAATGTTCCGGTTGCACTGAAA
GGCACCACCAATCAGAGCCGTCATACCAGCTATCAGCAGGGTATGTGTAAAGTTAGCTTTCAAGA
ACTGAACAACGAGGTGCTGTAACCTAGGCGCTTCAACGGAACGGATCTTACATATCGGGGGGGTA
GGGGTTTTTTGTCTCGGAGACCAAGTAGGGCATAATACGACTCACTATAGGGAGACTATGGATAA
AGGAGGTAAATAATGAAAATCCTGTTTAGCCCGATTGGTAATAGCGATCCGTGGCGTAATGATCGT
GATGGTGCAATGCTGCATATTGTGCGTCATTATAATCTGGATAAAGTGGTGCTGTATTTCACCCGT
ACCATTTGGGAAGGTAATGAAAATCGCAAAGGCCACAAAATCTATGAATGGGAGAAAATTATCCAG
ACCGTTAGCCCGAATACCGAAGTGGAAATTATCATTGAAAATGTGGATAACGCCCAGGATTACGAT
GTGTTCAAAGAGAAATTCCATAAATATCTGAAAATCATCGAAGATAGCTACGAGGATTGCGAAATT
ATTCTGAATGTTACCAGCGGTACACCGCAGATGGAAAGCACCCTGTGTCTGGAATATATTGTTTAC
CCGGAAAACAAAAAATGCGTTCAGGTTAGCACCCCGACCAAAGATAGCAATGCAGGTATTGAATA
TAGCAACCCGAAAGACAAAGTGGAAGAATTTGAAATCGTGAACGAAGTCGAGAAAAAAAGCGAAA
AACGCTGCAAAGAAATCAACATTCTGAGCTTTCGTGAAGCCATGATTCGTAGCCAGATTCTGGGTC
TGATTGATAACTATGATTATGAAGGTGCCCTGAATCTGGTGAGCAATCAGAAAAGTTTTCGCAATG
GTAAACTGCTGCGTAAAAAACTGCTGAGCCTGACCAAACAAATCAAAACCCATGAAGTTTTCCCGG
AAATCAACGAAAAATATCGTGATGACGCCCTGAAAAAATCCCTGTTTCATTATCTGCTGCTGAACAT
GCGTTATAATCGTCTGGATGTTGCAGAAACCCTGATTCGTGTTAAAAGCATTGCAGAGTTTATCCT
GAAAACCTACATCGAAATTCATTGGCCGACCCTGATTATTGAGAAAGATGGTAAACCGTATCTGAA
CGATGAAGATAATCTGTCCTTCGTGTACAAATACAACCTGCTGCTGGAAAAACGCAAACAGAATTT
TGATGTTAGCCGTATTCTGGGCCTGCCTGCATTTATTGATATTCTGACCATTCTGGAACCGAATAG
CCAGCTGCTGAAAGAAGTTAACGCAGTTAACGATATTAATGGCCTGCGTAATAGCATTGCCCATAA
CCTGGATACCCTGAACCTGGACAAAAATAAAAACTACAAAAAAATCATGCTGAGCGTGGAAGCCAT
CAAAAATATGCTGCACATTAGCTTCCCTGAAATCGAGGAAGAGGATTATAACTATTTTGAAGAGAA
AAACAAAGAATTTAAAGAACTGCTGTAACTCGAGAGGTTACAGCCTGCATAATGTAGCATAACCCC
TTGGGGCCTCTAAACGGGTCTTGAGGGGTTTTTTGTGCCTATAGTTTGAAGCAGAATCGAATTTCT
GCCATTCATCCGCTTATTATCACTTATTCAGGCGTAGCACCAGGCGTTTAAGGGCACCAATAACTG
CCTTACAAAAAACCCCTAGCCGCCCGATAAGAGCGGGCTAGGGGTTCGAGTAAAAAAAATTACGC
CCCGCCCTGCCACTCATCGCAGTACTGTTGTAATTCATTAAGCATTCTGCCGACATGGAAGCCATC
ACAGACGGCATGATGAACCTGAATCGCCAGCGGCATCAGCACCTTGTCGCCTTGCGTATAATATT
TGCCCATCGTGAAAACGGGGGCGAAGAAGTTGTCCATATTGGCCACGTTTAAATCAAAACTGGTG
AAACTCACCCAGGGATTGGCTGAAACGAAAAACATATTCTCAATAAACCCTTTAGGGAAATAGGCC
AGGTTTTCACCGTAACACGCCACATCTTGCGAATATATGTGTAGAAACTGCCGGAAATCGTCGTGG
TATTCACTCCAGAGCGATGAAAACGTTTCAGTTTGCTCATGGAAAACGGTGTAACAAGGGTGAACA
CTATCCCATATCACCAGCTCACCGTCTTTCATTGCCATACGGAACTCCGGGTGAGCATTCATCAGG
CGGGCAAGAATGTGAATAAAGGCCGGATAAAACTTGTGCTTATTTTTCTTTACGGTCTTTAAAAAG
GCCGTAATATCCAGCTGAACGGTCTGGTTATAGGTACATTGAGCAACTGACTGAAATGCCTCAAAA
TGTTCTTTACGATGCCATTGGGATATATCAACGGTGGTATATCCAGTGATTTTTTTCTCCATTTTAG
CTTCCTTAGCTCCTGAAAATCTCGATAACTCAAAAAATACGCCCGGTAGTGATCTTATTTCATTATG
GTGAAAGTTGGAACCTCTTACGTGCCGATCAAGGTCTCATTTTCGCCAAAAGTTGGCCCAGGGCT
TCCCGGTATCAACAGGGACACCAGGATTTATTTATTCTGCGAAGTGATCTTCCGTCACAGGTATTT
ATTCGGCGCAAAGTGCGTCGGGTGATGCTGCCAACTTACTGATTTAGTGTATGATGGTGTTTTTGA
GGTGCTCCAGTGGCTTCTGTTTCTATCAGCTGTCCCTCCTGTTCAGCTACTGACGGGGTGGTGCG
TAACGGCAAAAGCACCGCCGGACATCAGCGCTAGCGGAGTGTATACTGGCTTACTATGTTGGCAC
TGATGAGGGTGTCAGTGAAGTGCTTCATGTGGCAGGAGAAAAAAGGCTGCACCGGTGCGTCAGC
AGAATATGTGATACAGGATATATTCCGCTTCCTCGCTCACTGACTCGCTACGCTCGGTCGTTCGAC
TGCGGCGAGCGGAAATGGCTTACGAACGGGGCGGAGATTTCCTGGAAGATGCCAGGAAGATACT
TAACAGGGAAGTGAGAGGGCCGCGGCAAAGCCGTTTTTCCATAGGCTCCGCCCCCCTGACAAGC
ATCACGAAATCTGACGCTCAAATCAGTGGTGGCGAAACCCGACAGGACTATAAAGATACCAGGCG
TTTCCCCCTGGCGGCTCCCTCGTGCGCTCTCCTGTTCCTGCCTTTCGGTTTACCGGTGTCATTCC
GCTGTTATGGCCGCGTTTGTCTCATTCCACGCCTGACACTCAGTTCCGGGTAGGCAGTTCGCTCC
AAGCTGGACTGT

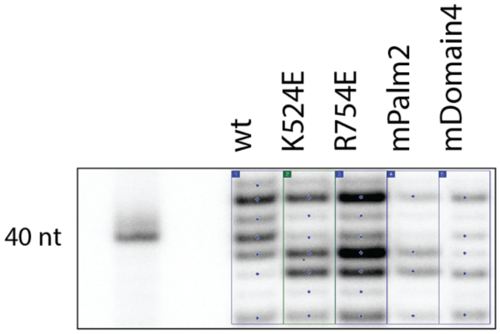


| | Amt. relative to wt |
|---|---|
| wt | 1.0 |
| K524E | 0.9 |
| R754E | 1.64 |
| mPalm2 | 0.43 |
| mDomain4 | 0.42 |

**Figure S3. Quantitation of crRNA associated with Cas10-Csm variants.** An image of a PAGE separation showing crRNA bands in the range of 36-43 nucleotides (nt). Blue dots in the image indicate bands that were quantitated by ImageQuant software. The sum of the amount of crRNA bands in each lane is presented in a table expressed in relative units.

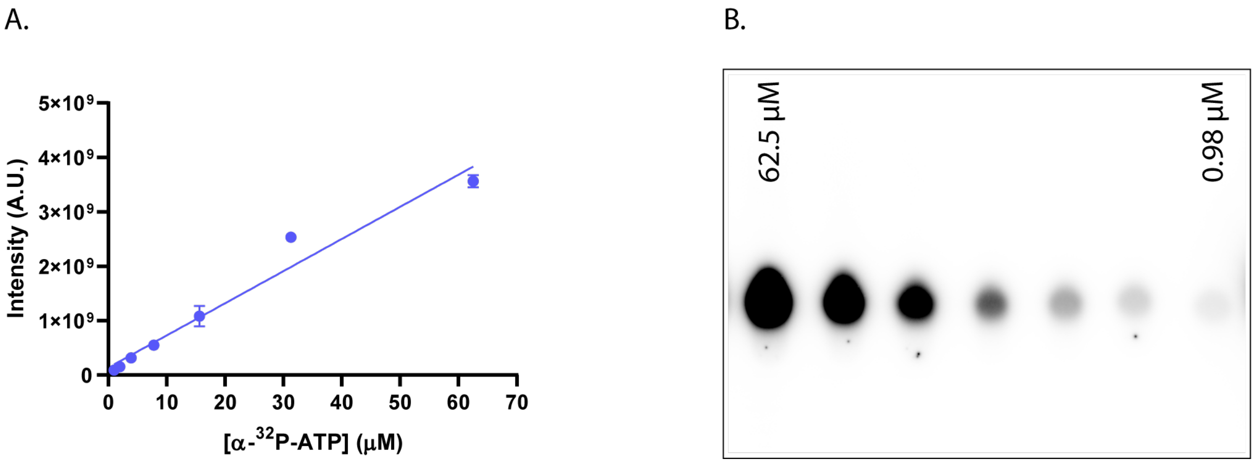


**Figure S4. A standard curve of α-$^{32}$P-ATP.** (A) Linear regression of the integrated intensities of α-$^{32}$P-ATP. (B) A representative TLC plate with serial dilutions of α-$^{32}$P-ATP.

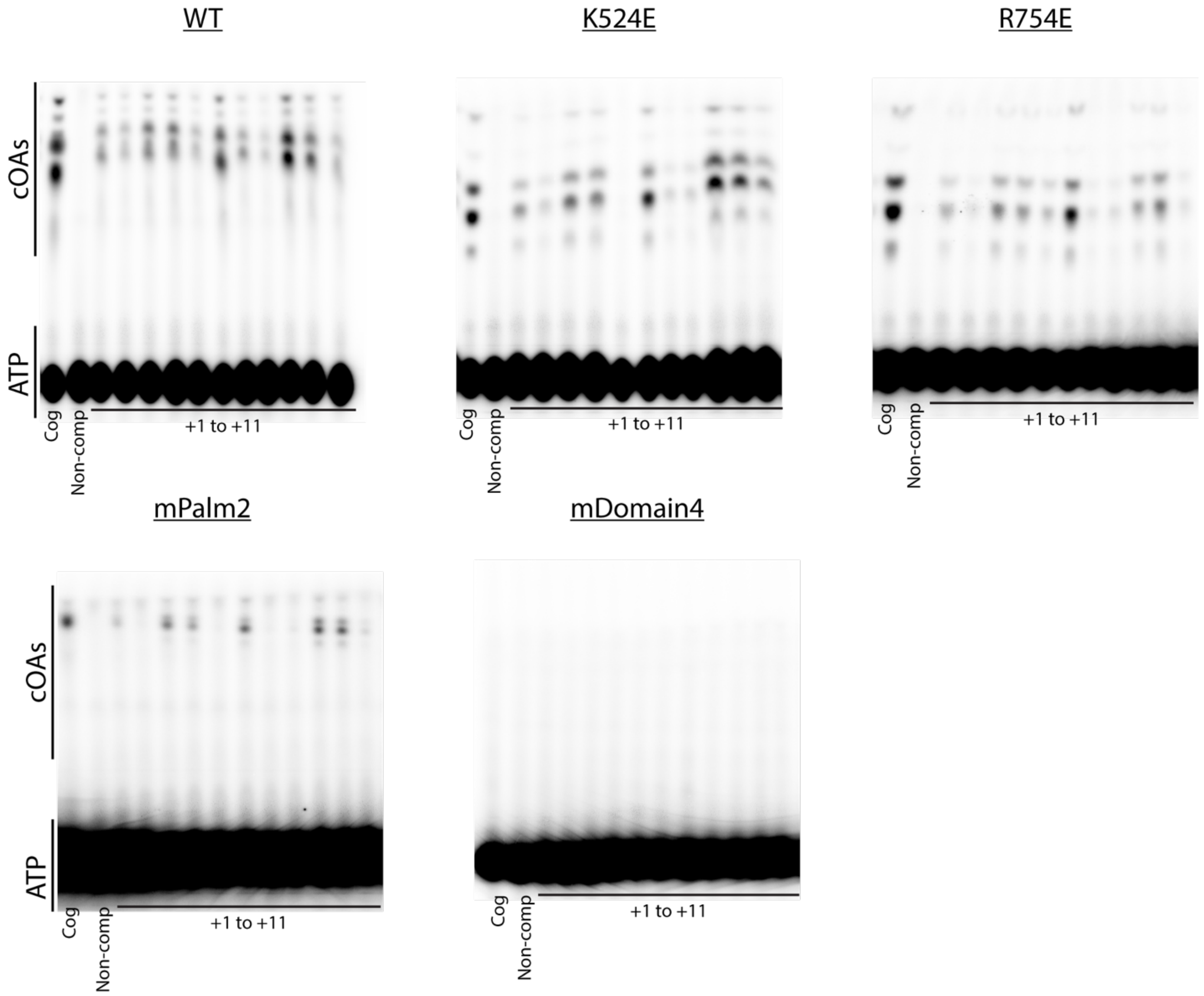


**Figure S5. Uncropped images of TLC plates reported in Figure 9.**

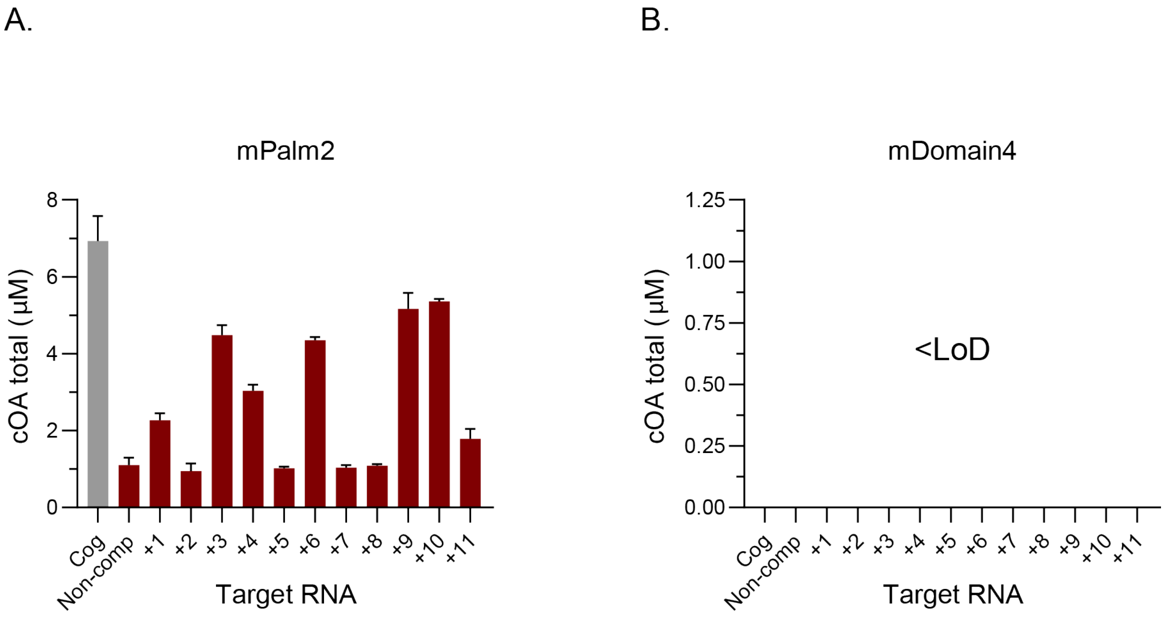


**Figure S6. COA synthesis by mPalm2 and mDomain4 Cas10-Csm variants.**
